# Spore type-specific gene expression profiles underlying development and leaf infection processes of *Colletotrichum graminicola*

**DOI:** 10.1101/2025.11.19.689217

**Authors:** Disha Rathi, Karsten Andresen, Rolf Daniel, Marco Alexandre Guerreiro, Matthias Kretschmer, James Kronstad, Minou Nowrousian, Stefanie Pöggeler, Anja Poehlein, Lars M Voll, Daniela Elisabeth Nordzieke

## Abstract

*Colletotrichum graminicola* causes significant losses of the staple crop maize worldwide. The fungus produces two distinct asexual spore types, oval and falcate conidia, which show unique processes in development and plant interaction. Based on genome resequencing of our laboratory strain (CgM2/ M1.001), we investigated the gene expression profiles of oval and falcate conidia during development and early leaf infection using RNA-seq. Our results reveal specific gene expression profiles between the two spore types, indicating fundamental differences in their developmental programs that reflect different modes of infection. We identified expression patterns discriminating both conidia types from mycelium and spore type-specific ones for genes encoding transcription factors, conserved fungal developmental genes, transporters, genes of secondary metabolite clusters, and pathogenicity-related functions, including effectors and carbohydrate-active enzymes (CAZymes). Our study shows that despite the identical genomic basis, oval and falcate conidia show unique transcriptomes across vegetative development and early plant interaction. Taking together, these results provide new insights into the molecular mechanisms determining the biology of *C. graminicola* and its interaction with the plant host.

## Introduction

*Colletotrichum* is a genus of phytopathogenic fungi within the *Glomerellaceae* family and poses a significant threat to global food security by causing devastating diseases in numerous crop species consumed by humans (Gomes et al. 2021). *Colletotrichum* species interact with a broad range of plant hosts, including cereals (maize, rice, sorghum), fruits (bananas, mango, avocado, citrus, strawberry), legumes (soybean, common bean), and vegetables, which makes it a major constraint in global food production (Baroncelli et al. 2017, Joshi 2018, Samarakoon et al. 2018, Damm et al. 2019, Gomes et al. 2021, Talhinhas and Baroncelli 2021, Boufleur et al. 2022, Gupta et al. 2022, Liu et al. 2022). Species of the *Colletotrichum* genus are grouped into 15 species complexes and exhibit a remarkable diversity in their infection strategies, ranging from biotrophs, necrotrophs, and hemibiotrophs to endophytes, allowing them to colonize and obtain nutrients from their hosts in various ways (Baroncelli et al. 2017, De Silva et al. 2017, Talhinhas and Baroncelli 2021). The genus has been ranked as the eighth most important group of plant pathogenic fungi globally, reflecting its significant scientific and economic impact (Cannon et al. 2012, Dean et al. 2012).

One of the most important *Colletotrichum* species infecting cereals is *C. graminicola*, which primarily infects maize (*Zea mays*) and causes substantial losses worldwide (Frey et al. 2011, Belisário et al. 2022). Maize is among the world’s most extensively cultivated crops, playing an essential role in global food systems as a fundamental source of nutrition for humans and animals and serving as a key raw material in various industrial applications including bioenergy (Kandil et al. 2020, Revilla et al. 2022, Assaf et al. 2024). *C. graminicola* can infect all plant parts and can be found throughout the growing season, making it a significant threat to maize production (Becerra et al. 2023). The disease manifests in anthracnose leaf blight (ALB), anthracnose stalk rot (ASR), and systemic plant infections, whereas current analyses estimate that ASR is the most important form causing 10-20% annual loss in maize harvest worldwide (Belisário et al. 2022). For efficient plant infection, *C. graminicola* produces two distinct spore types, oval and falcate conidia (Bergstrom and Nicholson 1999, Nordzieke et al. 2019b, Rudolph et al. 2025b). In contrast to other asexual spore type pairs from plant pathogenic fungi like *Magnaporthe oryzae*, oval and falcate conidia both contain the full set of organelles and are able to germinate under favorable conditions (Panaccione et al. 1989, Zhang et al. 2014). However, the conidiation process of both spore types differs significantly: falcate conidia develop from conidiophores in acervuli on infected leaves, whereas oval conidia are formed from hyphae in parenchyma cells in both leaf and stem lesions (Sukno et al. 2008, Nordzieke et al. 2019b, Belisário et al. 2022). Recent characterization of both conidia types revealed that they show significant differences in spore hydrophobicity, developmental processes and the adaptation to leaf or root infection (Chaky et al. 2001, Nordzieke et al. 2019b, Rudolph et al. 2025b). Newly generated falcate conidia have a pronounced dormant stage, which is controlled by the secondary metabolites mycosporine glutamine and mycosporine alanine (Leite 1992, Nordzieke et al. 2019b). Either exposure to glucose or attachment to hydrophobic surfaces on a polar adhesive strip induce a break of dormancy to give rise to either vegetative or pathogenic development, respectively (Leite 1992, Chaky et al. 2001, Vasselli et al. 2022). In contrast, oval conidia lack a dormant stage and start to germinate even under nutrient limited conditions (Nordzieke et al. 2019b). Oval conidia-derived germlings sense and grow towards each other by following chemical gradients to form a vegetative hyphal network by germling fusions, a process that has never been observed for falcate conidia so far (Nordzieke et al. 2019b, Rudolph et al. 2025a). The early developmental processes of both conidia types have consequences for the infection strategies on leaves: falcate conidia form melanized appressoria from short germ tubes which will subsequently form penetration pegs and infect the subjacent plant material. Oval conidia, however, form penetration structures from hyphae and germ tubes (hyphopodia) only when conidia are present in high spore densities. Thus, falcate conidia cause more severe symptoms on leaves when applied in lower spore numbers (Nordzieke et al. 2019b).

In the research community, the *C. graminicola* isolate CgM2 (M1.001), which was collected in Missouri from infected maize, is used for scientific investigations due to the reliable symptom development on leaves and stalks (Forgey et al. 1978). The first genome sequence of *C. graminicola* M1.001/CgM2 was generated in a community effort in 2012 using Sanger and 454 sequencing platforms, with a resulting reference assembly spanning 50.9 Mb. Due to the high number of repetitive sequences in the reference strains’ genome, the scaffolding was done based on the *Colletotrichum higginsianum* genome (O’Connell et al. 2012). A follow up study combined deep sequencing RNA-seq data with sophisticated bioinformatics strategies to improve genome annotation, which included identification of several hundred novel transcripts, improvement of gene models, and detection of candidate genes for alternative splicing (Schliebner et al. 2014). In 2023, a subsequent genome assembly of the same strain was generated using a combination of PacBio and Illumina sequencing technologies. Overall, 10 main chromosomes and 3 minichromosomes were identified, with a total genome size of 57.43 Mb, providing a high-quality genome sequence for this pathogen (Becerra et al. 2023). Those genomes were the bases of RNA-seq studies, focusing on different stages of leaf infection caused by falcate conidia. In those studies, genes encoding effectors, enzymes for secondary metabolites, transporters, hydrolases, and plant defense responses were identified as relevant for leaf infection by this spore type (O’Connell et al. 2012, Torres et al. 2016, Miranda et al. 2017).

In this study, we investigated gene expression profiles of oval and falcate conidia during development and early leaf infection via RNA-seq, aiming to explain the determining factors shaping spore type-specific behavior shown in previous work. To generate a sound basis for RNA-seq evaluation, we applied Nanopore and Illumina techniques to resequence the genome of our laboratory strain, which is present today in several lines around the globe, all originating from CgM2 (M1.001). We focused on the analysis of several functional gene groups relevant for development and pathogenicity, namely conserved fungal developmental genes and genes encoding for transcription factors, membrane transport proteins, secondary metabolite enzymes (encoded in gene clusters), effectors, and carbohydrate-active enzymes (CAZymes). Overall, we found significant overlaps and differences in gene expression profiles comparing oval conidia, falcate conidia and mycelium, indicating biological similarities and differences of the freshly harvested spores. Furthermore, we identified genes that encode pathogenicity relevant functions such as effectors, CAZymes, and transporters; some of these functions show spore type-specific expression patterns. The reported results support the view that oval and falcate conidia-derived hyphae have a specific identity which is maintained during development and early pathogenic interaction with the plant, shaping those processes in a spore type-specific way.

## Materials and Methods

### Strains, media and growth conditions

The wild-type strain CgM2 (M1.001) of *C. graminicola* (Ces.) G.W.Wilson was provided by R. L. Nicholson to H. Deising (Martin-Luther University Halle-Wittenberg), who provided it to our laboratory. CgM2 was renamed to M1.001 by B. Hanau. Currently, both names are used as synonyms for the same strain (Forgey et al. 1978, Werner et al. 2007, O’Connell et al. 2012, Aliyeva-Schnorr et al. 2025). Falcate conidia were generated by growing *C. graminicola* on oatmeal agar (OMA, 1l: 50 g ground oat flakes (Alnatura, Darmstadt, Germany)) for 14-21 d. To stimulate falcate conidia generation, *C. graminicola* was exposed to red (660–665 nm) and blue (450–455 nm) light in a ratio of 3:1 at 23°C. For the generation of oval conidia, mycelial plugs or a falcate conidia spore solution were inoculated in liquid complete medium with 0.5 M of sucrose (CMS, 1l: 55.5 mM glucose, 0.1% yeast extract, 0.1% peptone, 0.5 M sucrose, 5.3 mM Ca(NO_3_)_2_) 0.073 mM KH_2_PO_4_, 1.04 mM MgSO_4_, 0.46 mM NaCl) and shaken (80 g, 23°C) in darkness for two days. After shaking, the cultivation was continued for 3–5 days in darkness without agitation (23°C; Rudolph et al. 2025b). Mycelium was grown in liquid CM medium (CM, 1l: 1% glucose, 0.1% yeast extract, 0.1% peptone, 5.3 mM Ca(NO_3_)_2_), 0.073 mM KH_2_PO_4_, 1.04 mM MgSO_4_, 0.46 mM NaCl) using either mycelial plugs or spore solutions of the CgM2 wild-type strain, followed by cultivation for 5 d at 23°C.

### Preparation of genomic DNA for genome resequencing

Genomic DNA (gDNA) was prepared from mycelium grown for 5 d in liquid CM medium. To allow the sequencing of full chromosomes of CgM2, the NucleoBond^®^ HMW DNA kit (Macherey-Nagel, Düren, Germany; ref 740160.2) was used for high molecular weight gDNA extraction according to the manufacturer’s instructions.

Concentration of the isolated DNA was determined using the Qubit^®^ dsDNA HS Assay Kit as recommended by the manufacturer (Life Technologies GmbH, Darmstadt, Germany). Shotgun libraries were prepared using the Illumina^®^ DNA Prep, (M) Tagmentation kit and Nextera™ DNA CD Indexe kit (96 Indexes) as recommended by the manufacturer (Illumina Inc., San Diego, CA, USA). To assess quality and size of the libraries, samples were run on an Agilent Bioanalyzer 2100 using an Agilent High Sensitivity DNA Kit as recommended by the manufacturer (Agilent Technologies, Waldbronn, Germany). Concentration of the libraries were determined using the Qubit^®^ dsDNA HS Assay Kit as recommended by the manufacturer (Life Technologies GmbH, Darmstadt, Germany). Sequencing was performed by using the NovaSeq6000 instrument (Illumina Inc., San Diego, CA, USA) using the NovaSeq6000 SP Reagent Kit (v1.5) and the NovaSeq XP 2-Lane Kit (v1.5) for sequencing in the paired-end mode 2x250 cycles.

For Nanopore sequencing 1.2 µg of high molecular weight DNA (HWD) was used for library preparation using the Ligation Sequencing Kit 1D (SQK-LSK109) as recommended by the manufacturer. Sequencing was performed for 72 h using a MinION device Mk1B and a SpotON Flow Cell R9.4.1 as recommended by the manufacturer (Oxford Nanopore Technologies) using MinKNOW software (22.05.5) for sequencing and Guppy (v7.1.3) in high accuracy mode for basecalling.

### Preparation of RNA samples and RNA extraction procedures

RNA of mycelium and freshly harvested oval and falcate conidia of wild-type CgM2 were isolated and prepared for subsequent sequencing. CgM2 falcate conidia were harvested after 28 d cultivation on OMA using 0.02% Tween 20 solution. Oval conidia were generated in liquid CMS medium and separated from mycelium by filtration (Miracloth, EMD Milipore Corp, Burlington, USA; ref 475855-1R). Both spore types were centrifuged (4,000 g, 10 min), the supernatant discarded and the pellet collected on a sterile filter disc (grade 3hw, Sartorius, Göttingen, Germany; LOT 10-038) with the help of a vacuum pump (Type: PM12640-026.3, Nr. 02128309, Biometra, Germany).

To assess gene expression during spore germination and germling fusion, distinct harvesting time points were chosen to reflect differences in conidia-specific developments. Oval conidia rapidly germinate also without the requirement of nutrient sources and reaches levels of more than 80% germination after 6 h. Germling network formation by oval conidia starts at that timepoint but reaches its maximum after about 17 h post inoculation (Nordzieke et al. 2019b). Germination of falcate conidia, however, is tightly controlled and several factors ensure spore dormancy und nutrient limiting conditions (Leite and Nicholson 1992, Chaky et al. 2001, Vasselli et al. 2022). To identify genes responsible for the different progression of vegetative spore development, we first adapted the cultivation conditions to induce the desired developmental processes and allowed at the same time for germling harvest and RNA extraction. We found that inoculation of 50 µl of conidia diluted in 450 µl of distilled water together with 500 µl 50 mM NaNO_3_ solution (= 1 ml total volume) in 6 well plates (TC-Platte 6 Well Cell+,F; Sarstedt, Germany, ref 83.3920.300), which additionally contained pieces of miracloth (Miracloth, EMD Milipore Corp, Burlington, USA; ref 475855-1R) provided the required conditions to mimic development as observed previously on water agar (Nordzieke et al. 2019b). To ensure further oval conidia germination and germling fusion rates or dormancy of falcate conidia as observed previously, spore titers were adjusted for both spore types (falcate conidia (c = 2.1 × 10^7^ml^-1^) or oval conidia (c = 2.6 × 10^7^ml^-1^). After incubation at 23°C, liquid and fungal tissue were sampled by pipetting on a filter disc (grade 3hw, Sartorius, Göttingen, Germany; LOT 10-038) with the help of a vacuum pump (Type: PM12640-026.3, Nr. 02128309, Biometra, Germany). The dried filter was frozen in liquid nitrogen until RNA was extracted.

In a previous study, we found that oval and falcate conidia show a different behavior prior to leaf infection (Nordzieke et al. 2019b). Falcate conidia form appressoria from short germ tubes, followed by penetration of the plant surface. Oval conidia form networks by germling fusions when present in high spore densities. At the same time, hyphopodia, melanized penetration structures from hyphae, emerge and penetrate the plant surface (Nordzieke et al. 2019b). Although germling fusion and hyphopodia formation are spore concentration-dependent processes, network formation is not a prerequisite for the development of hyphopodia (Nordzieke 2022). To rule out this spore-density dependent factor when comparing transcriptomes of falcate and oval conidia early leaf infections, we choose spore titers enabling the generation of likewise symptoms strengths in both conidia types (Nordzieke et al. 2019b). To assess which genes are regulating the spore type-specific leaf infection strategies, we prepared RNA from infected maize leaves (cultivar Mikado, KWS SAAT SE, Einbeck, Germany). *Zea mays* plants were grown in a 4:1 mix of soil (SP Vermehrung, Einheiterde Werkverband e.V., Sinntal-Altengronau, Germany, Artikel-Nr. 11-01500-40) and vermiculite (grain size 2–3 mm, Isola Vermiculite GmbH, Sprockhövel, Germany) in a controlled environment (PK 520 WLED plant chamber (Poly Klima Climatic Growth System, Freising, Germany), day/ night cycle of 12 h (26°C /18°C)) for 16 d. Secondary leaves of the grown plants were placed in square petri dishes (Petri Dish 100x100x20mm, Sarstedt, 16 Park Way, Mawson Lakes, South Australia 5095, REF 82.9923.422) on top of wet blotting paper (BF2 580 x 600 mm, Sartorius, Göttingen Germany). Conidia suspensions were prepared in 0.01% Tween solutions to ensure attachment to the leaf surface (Gu et al. 2024). Then 10-12 drops, each of 10 µl c = 10^5^ ml^-1^), were applied to the leaves. The square plates were sealed to ensure high humidity. After incubation for 24 h at 23°C, infection spots were collected (4mm Miltex Biopsy Punch with Plunger, Integra, Mansfield, USA; ref 33-34-P/25) in an Eppendorf tube and stored in liquid nitrogen until RNA extraction.

RNA extraction was performed using a RNeasy Plant mini kit (Qiagen, Hilden, Germany, ref 74904). The eluted RNA was treated with DNase (40 μg) using Turbo DNA free kit (Invitrogen, Vilnius, Lithuania, ref AM1907) to remove DNA contamination. Clean up of the samples was performed using a RNeasy mini elute clean up kit (Qiagen, Hilden, Germany, ref 74204). Depending on the follow up application, RNA was either prepared for RNA sequencing or transcribed into complementary DNA (cDNA) as template for qRT-PCR (Figure A File S1).

### Validation of RNA-seq Data by quantitative Real-Time PCR (qRT-PCR) analysis

For quantitative PCR, cDNA samples were prepared using purified RNA samples and an iScript cDNA synthesis kit (BIO RAD, CAT-1708891). qRT-PCR was performed using SsoAdvanced universal SYBR green supermix (BIO RAD, CAT-1725271). For detection, a CFX Connect Real Time PCR detection system (Bio-Rad Laboratories, Hercules, CA, USA) was used. The GAPDH gene was used as the internal reference (housekeeping gene). Relative expression levels of target genes were calculated using the ΔΔCt method (Livak and Schmittgen 2001), normalizing each sample to its corresponding GAPDH value. Fold change values (log2 fold change ≤ –1 or ≥ 1, a value of -1 means that a gene is two-fold down-regulated, whereas a value of 1 means that the gene is two-fold up-regulated in this comparison) were applied for comparative analysis. Heatmaps representing the expression profiles from both RNA-seq and qRT-PCR Datasets were generated using the iDEP.96 platform (Ge et al. 2018).

### Genome assembly and annotation

Oxford Nanopore reads of at least 10 kb were assembled with Canu v2.2 (Koren et al. 2017) with parameter genome size = 58 m. The resulting assembly was corrected with four rounds of Racon v1.4.3 (Vaser et al. 2017) using the Oxford Nanopore reads and four rounds of Pilon v1.24 (Walker et al. 2014) using the trimmed Illumina reads. The resulting 17 contigs were compared with the previously published genome V1 including the published optical map data (O’Connell et al. 2012) to assign the contigs to chromosomes resulting in 13 contigs representing chromosomes. Racon and Pilon with the Nanopore and Illumina reads, respectively, were used to close the gaps introduced in the scaffolding. The resulting gapless contigs were analyzed for the presence of telomeric repeats (sequence TTAGGG) using a custom-made Perl script. Telomeric repeats were found at both ends of all contigs except for contig 10 that has a telomeric repeat at one end and 24 copies of the ribosomal RNA (rRNA) genes at the other end (most likely preventing assembly of telomeric repeats at that end). BUSCO (Benchmarking Universal Single-Copy Orthologs v.5.2.2) analysis (Manni, Berkeley et al. 2021) with BUSCO Dataset fungi odb10 showed 97.8 % completeness.

For gene annotation, RNA-seq reads derived from the combined RNA of vegetative mycelium and two different spore types were assembled with Trinity v2.15.1 (Grabherr et al. 2011). The resulting *de novo* assembled transcripts were used together with the predicted proteins from the published V1 genome (O’Connell et al. 2012) for genome annotation with MAKER (v3.01.03; Cantarel et al. 2008) as well as for the BRAKER2 pipeline (v2.1.2; Stanke et al. 2006, Stanke et al. 2008, Li et al. 2009, Hoff et al. 2016, Hoff et al. 2019, Brůna et al. 2021). For the latter, two separate runs were conducted with protein and transcriptome input, respectively, and the resulting output files were combined using TSEBRA. The gene models from MAKER and BRAKER2 were combined using custom-made Perl scripts. Subsequently, another MAKER run was conducted using the predicted proteins and RNAs from the combined MAKER and BRAKER2 analyses together with predicted proteins from a recently published *C. graminicola* genome assembly (Becerra et al. 2023). BUSCO (Benchmarking Universal Single-Copy Orthologs v.5.2.2 (Manni et al. 2021), analysis of the resulting predicted proteins with the fungi_odb10 Dataset showed 97.1 % completeness.

### Annotation of relevant gene groups

#### Transcription factor encoding genes

Data from the 312 predicted transcription factor genes identified previously in *Neurospora crassa* OR74A (Carrillo et al. 2017) was used in this study. The predicted genome of *C. graminicola* was aligned with these predicted TFs using local BLAST+ command line homology searches. The matches of the discontiguous MegaBLAST were further scrutinized by a BLASTN (BLAST+ 2.15.0 version) search run against the predicted genome in order to remove TF-unrelated as well as mispredicted sequences. The final list contains all the transcription factors with E value of <1e^-05^.

#### Effector encoding genes

Locus Tags of effector encoding genes were assigned based on previous effector gene identification performed by Becerra and coworkers (Becerra et al. 2023).

#### Genes encoding membrane transporters

Genes encoding membrane transport proteins were identified by a multilayered approach. Genes that had already been annotated in the genome version v4.0 (O’Connell et al. 2012) were identified by BLAST searches. The remaining genes were filtered by WoLF PSORT (https://wolfpsort.hgc.jp/) for plasma membrane, ER, vacuolar or mitochondrial localization and after that, by PFAM domain predictions typical for transport proteins (e.g. MFS_1, Sugar_tr, AA_permease, etc.). In this way, 104 novel transporter genes were identified. These candidate genes were then annotated by BLAST (version 2.14.0) searches against Transporter Classification Database (https://www.tcdb.org), the *Saccharomyces* Genome Database (https://www.yeastgenome.org) and the NCBI Reference Sequences database (https://www.ncbi.nlm.nih.gov), as described by (O’Connell et al. 2012).

#### Genes encoding CAZymes

The predicted proteome of *C. graminicola* was scanned against the dbCAN3 v11 database (Zheng et al. 2023) for CAZymes by using all tools (HMM, DIAMOND, dbCAN-sub) within the run_dbcan4 tool. Matches with an HMM e-value below 1e^-15^ and coverage greater than 0.35 were considered for further steps. Additionally, the proteome was screened by the CUPP online server (Barrett et al. 2020a) with default settings and considering matches with a significance score above 5. Matches identified exclusively by DIAMOND were excluded from further analyses. Protein classification was based on consistent matches across all used tools, while inconsistent matches were disregarded for subsequent CAZyme analyses. Substrate prediction was achieved by using the dbCAN_sub database (Zheng et al. 2023).

#### Genes encoding regulators of development

For identification of autophagy related proteins, proteins involved in signaling (MAP kinase pathways, NOX proteins, STRIPAK components, pheromone-signaling proteins), transcription factors involved in developmental processes and melanin biosynthesis proteins, a reciprocal BLASTP (version 2.0.10) and a BLASTN (BLAST+ 2.15.0 version) analysis was conducted based on *Sordaria macrospora* and *Aspergillus flavus* proteins sequences, respectively. The highest sequence identity and e-value threshold <1e^-05^ and a continuous overlap of 50% over the query sequence was used for the detection of *C. graminicola* homologs.

#### Secondary metabolite biosynthesis genes

In order to locate secondary metabolite biosynthesis gene clusters, the genome was analyzed with antiSMASH version fungiSMASH 7.0.0 (Blin et al. 2021).

#### RNA-seq analysis

RNA extraction was performed using a RNeasy Plant mini kit (Qiagen, Hilden, Germany, ref 74904). The eluted RNA was treated with DNase (40 μg) using Turbo DNA free kit (Invitrogen, Vilnius, Lithuania, ref AM1907) to remove DNA contamination. RNA was prepared for RNA sequencing by clean-up of the samples using a RNeasy mini elute clean up kit (Qiagen, Hilden, Germany, ref 74204).

The NEB Next Poly(A) mRNA Magnetic Isolation Module (New England BioLabs, Frankfurt am Main, Germany) was used to reduce the amount of rRNA-derived sequences. For sequencing, the strand-specific cDNA libraries were constructed with a NEB Next Ultra II Directional RNA library preparation kit for Illumina and the NEB Next Multiplex Oligos for Illumina (96) (New England BioLabs, Frankfurt am Main, Germany). To assess quality and size of the libraries samples were run on an Agilent Bioanalyzer 2100 using an Agilent High Sensitivity DNA Kit as recommended by the manufacturer (Agilent Technologies). Concentration of the libraries was determined using the Qubit® dsDNA HS Assay Kit as recommended by the manufacturer (Life Technologies GmbH, Darmstadt, Germany). Sequencing was performed on the NovaSeq 6000 instrument (Illumina Inc., San Diego, CA, USA) using NovaSeq 6000 SP Reagent Kit (100 cycles) and the NovaSeq XP 2-Lane Kit v1.5 for sequencing in the paired-end mode and running 2x 61 cycles.

RNA from 9 different samples with biologically independent triplicates per sample was analyzed by paired-end Illumina sequencing (Table A File S1). Reads were mapped to the genome sequence with Hisat2 v2.2.1 (Kim et al. 2019) allowing multi-mapping with default parameters and parameters for intron lengths set to --min-intronlen 20 and -- max-intronlen 4000. Reads mapping to annotated features were counted as described by (Teichert et al. 2012) with the modification that reads were strand-specific and counted if they mapped to the corresponding strand of the feature. Quantitative analysis of gene expression was done in R (v4.1.2; Team 2024) with DESeq2 1.34.0 (Love et al. 2014) using independent Filtering in the results function. Genes were identified as differentially expressed if they had an adjusted P-value (Padj) of <0.05 and log2 values of fold expression changes were either ≥1 for up-regulated or ≤1 for down-regulated genes. Venn diagrams of genes that were up- or down-regulated in different combinations were generated by identifying the corresponding genes using custom-made perl scripts (File S2), downstream processing using sorting functions in Excel (Microsoft) and generating of the diagram in Adobe Illustrator. The top 100 variable genes were identified using log2 fold change values and were analyzed using iDEP.96 (Ge et al. 2018) where log2 fold change <=-1 or >=1, padj <0.05 was used. Hierarchical clustering was performed based on correlation distance with average linkage. Z-score normalization (centered by subtracting the mean) was applied, and a cut-off Z-score of 4 was used for analysis. The generation of volcano blots was performed using the database VolcaNoseR (https://huygens.science.uva.nl/VolcaNoseR/; Goedhart and Luijsterburg 2020). VolcaNoseR settings were as follows: fold change threshold: -2 to 2; significant threshold: 2; use thresholds to annotate: changed and significant; criterion for ranking hits: Manhattan distance. Pathway enrichment analysis was performed using the GAGE method implemented in iDEP.96 (log2 fold change <=-1 or >=1, padj <0.05). Gene sets ranging from 5 to 2000 genes were included. Pathways were considered significant at an FDR cutoff of 0.05. Prior to enrichment analysis, genes with FDR ≥ 1 were excluded.

## Results

### Genome resequencing of the *C. graminicola* laboratory strain CgM2 (M1.001)

Since its collection, the CgM2/M1.001 isolate is used in several labs around the world. We therefore performed resequencing of our laboratory strain in this study to generate a reliable basis for data evaluation and interpretation. In this genome version V5, we obtained a gapless assembly with telomeric repeats at both ends of most chromosomes apart from chromosome 10, which contains rDNA repeats. In total, we assembled 13 chromosomes for *C. graminicola*, including the minichromosomes Chr11, Chr12 and Chr13. All chromosomes sum up to 57,600,233 base pairs in total. Using RNA-seq of a mixed sample containing biological material of *C. graminicola* mycelium, oval, and falcate conidia, 15,481 protein-encoding genes were annotated (Dataset S1). Although chromosome 1 is the longest, spanning 7,645,418 base pairs with 2,013 predicted genes, chromosome 2 contains the highest number of predicted genes (2,289). In total, our genome version V5 is very similar to V4 (Becerra et al., 2023), however, the total number of encoded proteins in V5 is increased by 400 (Table 1, Figure 1), most likely because different input data and algorithms were used for annotation.

**Figure 1:**
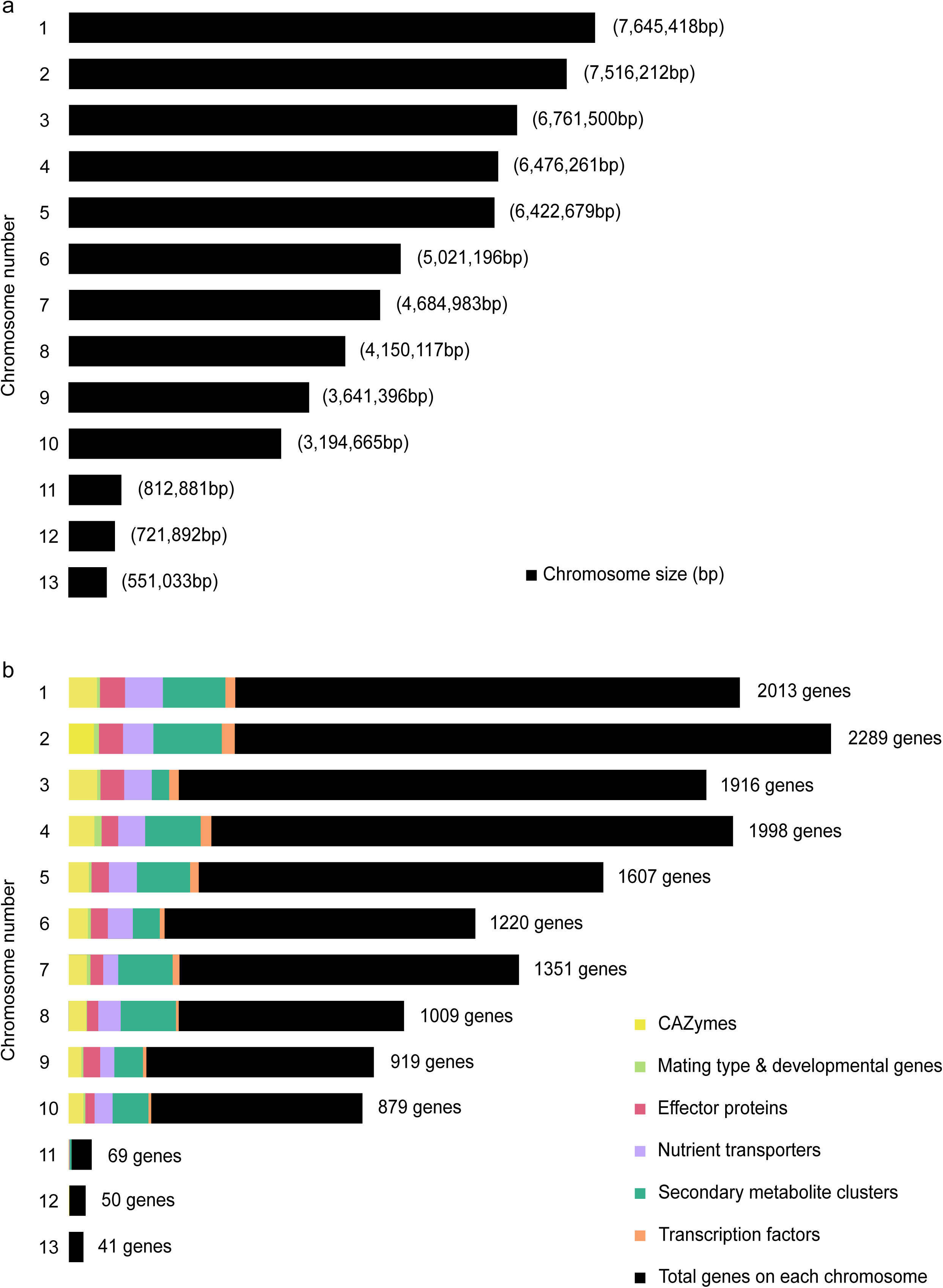
Overview of the *C. graminicola* assembly V5. (a) Size of different chromosomes in base pairs (bp). (b) Annotation of different functional gene groups based on the number of genes on 13 chromosomes, colored bars represent different functional categories. The black bar represents the total number of genes on one chromosome, including colored functionally categorized genes. Alt text: Bar charts summarizing the *C. graminicola* V5 genome assembly, with subfigures labelled from a to b, illustrating chromosome length and encoded genes.

**Table 1:** Comparison of assembly statistics for the *C. graminicola* M1.001 genome.

| Description | V1 | V4 | V5 |
| --- | --- | --- | --- |
| No. of contigs | 1,151 | 13 | 13 |
| Size (Mb) | 50.9 | 57.43 | 57.60 |
| Number of protein coding genes | 12,006 | 15,118 | 15,481 |
| GC content (%) | 49.12 | 45.98 | 46 |

V1 (O'Connell et al. 2012), V4 (Becerra et al. 2023), and V5, which was generated in this study applying a combination of Nanopore and Illumina sequencing techniques.

### Differential gene expression dynamics during spore development and maize leaf infection

In the 1970s, it was discovered that *C. graminicola* generates two distinct spore types (Nishihara 1975). Recent investigations from our group provided evidence that the differences culminate in specialization of the two spore types for the infection of different plant tissues (Nordzieke et al. 2019b, Rudolph et al. 2025b). In this study, we performed a comparative RNA-seq analysis to unravel the genetic basis for the observed biological differences between the spore types, focusing on three relevant timepoints identified in earlier work: cultivation under nutrient depletion conditions for 5 h and 16 h induce different developmental programs. In oval conidia, germination is induced after 5 h, which is followed by the generation of germling fusion, an active process at 16 h. Falcate conidia, however, remain dormant during both time points (Nordzieke et al. 2019b). In addition to the vegetative development, we included early leaf infection (1 dpi) samples. At this stage, high percentages of falcate conidia have formed penetrating appressoria on short germ tubes. Oval conidia, on the other hand, did germinate on leaves followed by the formation of germling fusions and penetrating hyphopodia. Originating from appressoria and hyphopodia alike, plant invasion has started and biotrophic primary hyphae are visible *in planta* (Nordzieke et al. 2019b, Nordzieke 2022). As controls, we included samples of freshly harvested oval and falcate conidia as well as mycelium of *C. graminicola*.

A Principal Component Analysis (PCA) was conducted to assess the quality of the data generated, and to visualize sample clustering and variation among the biological replicates. In our study, PC1 explains 44% of the total variance, while PC2 accounts for an additional 20%, indicating that these two components together capture a substantial 64% of the transcriptomic variation (Figure 2a). Biological replicates of the different samples clearly cluster according to the two spore types (falcate and oval conidia), and leaf infection (1dpi) with mycelium as an outgroup. Notably, falcate conidia (Fc) and oval conidia (Oc) samples were separated by PC1, demonstrating strong transcriptional differences between the two spore types. Within each group, samples taken during early vegetative development (5h, 16h) separate clearly, suggesting stage-specific expression changes. Mycelium samples (Myc) cluster apart from all conidial stages, emphasizing a specific expression profile during vegetative growth. The separation of leaf infection samples (Fc1dLF, Oc1dLF) from other samples predominantly by PC2 further supports the presence of transcriptional reprogramming during host-pathogen interaction.

**Figure 2:**
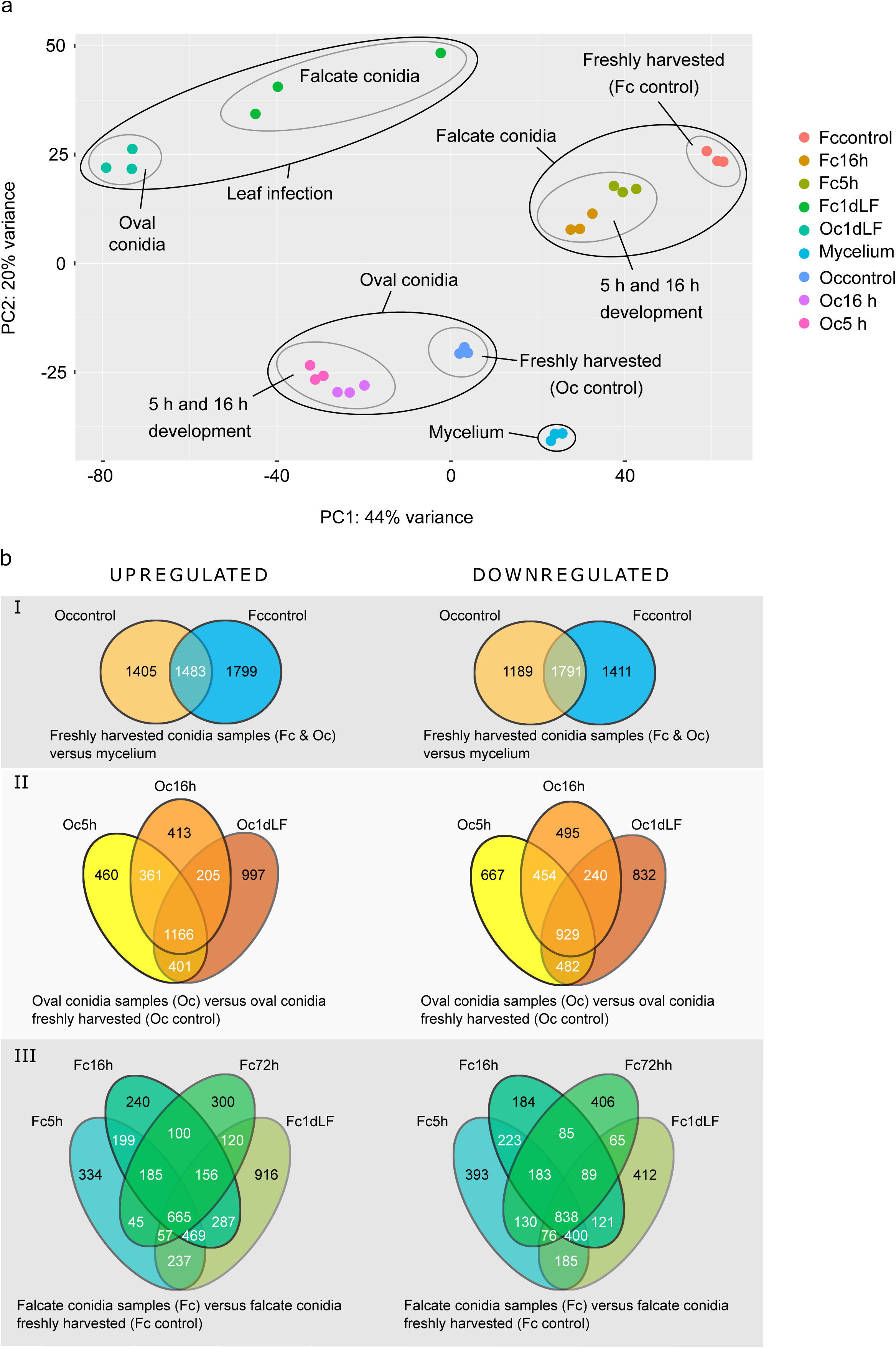
Differentially expressed genes in two different conidia types identified by DESeq2. Genes were identified as differentially expressed if they had an adjusted P-value (Padj) of <0.05 and log2 values of fold expression changes were either ≥1 for up-regulated or ≤1 for down-regulated genes. (a) Principal component analysis of RNA-seq data for all samples. The first two principal components explain 64% of variance. The four major groups of samples consist of mycelial samples, oval conidia (w/o the infection samples), falcate conidia (w/o the infection samples), and leaf infection samples. Oval conidia are displayed to improve visibility, they do not have statistical significance. (b) The Venn diagrams present an overview of differentially regulated genes of (I) freshly harvested falcate and oval conidia, (II, III) developmental stages of oval conidia (II) and falcate conidia (III). Alt text: Differentially expressed identified by DESeq2 using padj < 0.05 and log2 fold change ≥ 1, with subfigures labelled from a to b, illustrating a Principal Component Analysis (PCA) and Venn diagrams showing gene numbers of differentially expressed genes.

Visualization of the overall up and down-regulated genes in Venn Diagrams was used to explore differential gene expression during spore development and leaf infection (Figure 2b). Depending on the sample analyzed, we selected different reference samples for our entire analysis. Freshly harvested oval and falcate conidia were compared to mycelium samples to analyze whether those morphological stages differ in their transcriptome. For all stages developing from freshly harvested conidia (5 h, 16 h, 1 d leaf infection), the corresponding freshly harvested conidia sample served as control. To reflect probable differences in between early leaf infections of both conidia types, DEGs in between infection samples of oval and falcate conidia were identified. In freshly harvested oval and falcate conidia, >1000 genes were differentially regulated relative to vegetative mycelium and shared among both spore types, indicating that oval and falcate conidia are intrinsically diverse from vegetative hyphae. At the same time, >1000 genes are regulated in a spore type-specific manner relative to vegetative mycelium, underpinning the different nature of those conidia types compared to each other and reflecting the results obtained by PCA analysis. Furthermore, we observed differences in gene expression between the two conidia types during developmental transitions, 5 h and 16 h inoculations under nutrient-limiting conditions, and early leaf infection (1 dpi). In falcate conidia, 665 genes were differentially up-regulated and 838 differentially down-regulated across all developmental stages and during early leaf infection. Oval conidia displayed even stronger transcriptional dynamics, with 1,166 genes up-regulated and 929 down-regulated, reflecting a more pronounced transcriptional reprogramming associated with developmental progression and pathogenesis.

To obtain additional insight, we examined the 100 most variable expressed genes across developmental transitions, including freshly harvested conidia (control), critical timepoints for vegetative development (5 h, 16 h), and maize leaf infection (1 dpi, Figures B-E File S1). Pathway enrichment analysis performed with iDEP.96 revealed distinct expression signatures of freshly harvested oval and falcate conidia and early infection stages (Table B File S1). Specifically, genes involved in rRNA processing, cellular responses to DNA damage, and general stress responses were predominantly up-regulated in freshly harvested conidia. In contrast, during early leaf infection (1 dpi), there was significant enrichment for pathways linked to secondary metabolite biosynthesis, carbohydrate metabolism, and hydrolase activity.

### Validation of differential gene expression by qRT-PCR

To confirm the transcriptomic data obtained from RNA-seq (Dataset S2), a subset of differentially expressed genes was validated using quantitative real-time PCR (qRT-PCR, Figure 3, Table C File S1). The selection was based on the significance of the RNA-seq data (top 20 up- and down-regulated genes) and interesting regulation patterns dependent on spore type, developmental stage or early infection processes. Among the selected seven genes, COGRA5_13220 (Emopamil binding protein), COGRA5_06545 (Cutinase), COGRA5_00514 (Apoplastic effector) and COGRA5_09579 (Fungal RiPP-like secondary metabolite) showed interesting expression patterns, indicating a probable role of these in specific developmental transitions, such as dormancy (COGRA5_13220), dormancy breaking of falcate conidia (COGRA5_06545), germination induction of oval conidia (COGRA5_00514), and adhesion (COGRA5_09579). Regarding early leaf infection, we selected COGRA5_00442 (Carboxylesterase), which is overexpressed in falcate conidia during maize leaf infection, while COGRA5_14555 (Taurine catabolism dioxygenase TauD) showed increased expression in oval conidia under the same condition (Dataset S3). In contrast, COGRA5_09579 (Fungal RiPP-like secondary metabolite) is up-regulated in both spore types during infection, indicating shared pathogenicity-related functions. Overall, expression profiles obtained from qRT-PCR (Figure F File S1) were like the RNA-seq results and thus confirmed the robustness of the RNA-seq findings.

**Fig. 3:**
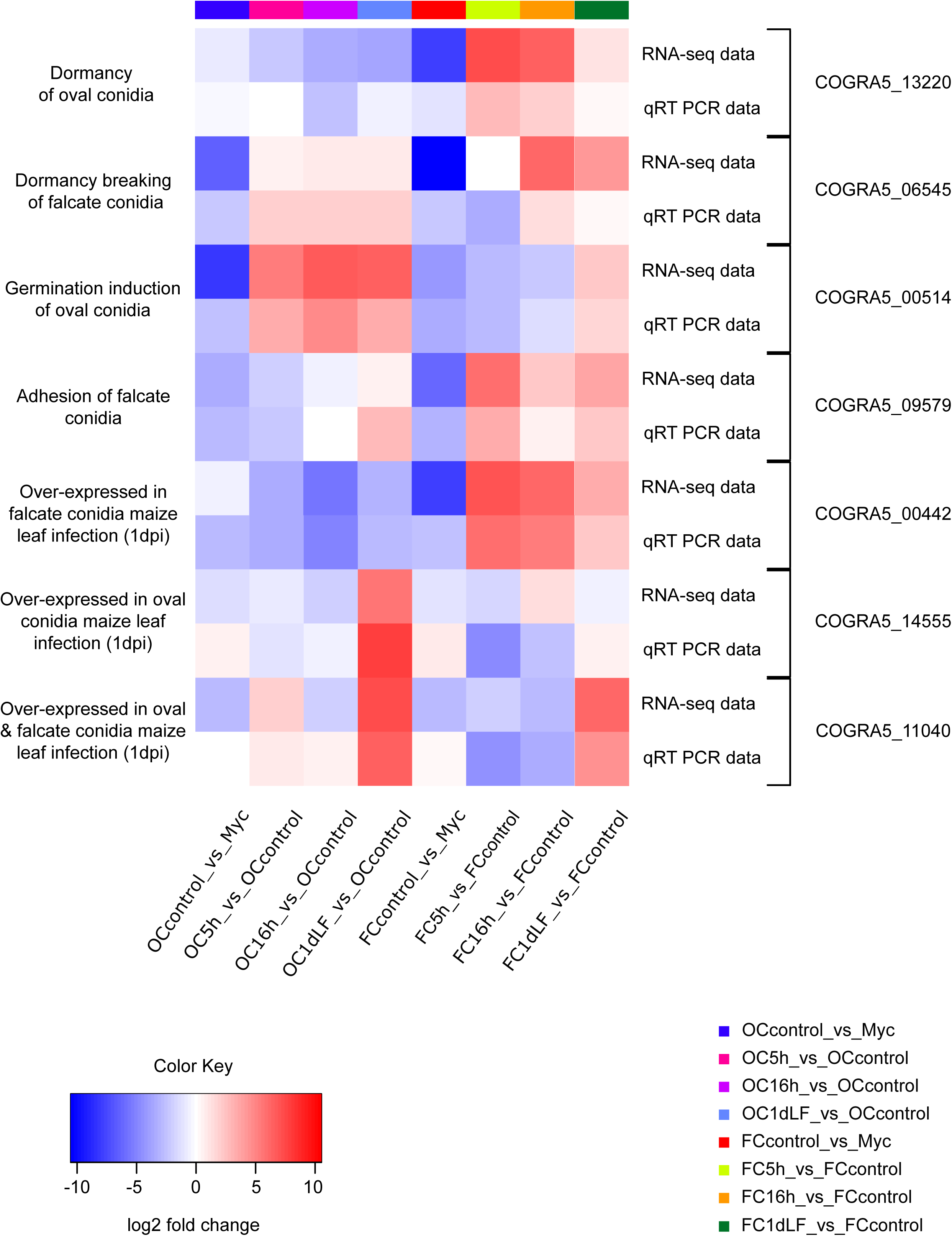
Comparative analysis of transcriptomic expression using both qRT-PCR and RNA-seq of selected genes. The data reflects the fold change correlations between these two methods, presented as a heat map for selected genes with a potential role in distinct developmental stages of falcate and oval conidia indicated (log2 fold change <=-1 or >=1, padj <0.05). The heatmap was created using iDEP.96 (Ge et al. 2018). Hierarchical clustering was performed based on correlation distance with average linkage. Z-score normalization (centered by subtracting the mean) was applied, and a cut-off Z-score of 4 was used for visualization. Red color depicts the up-regulated genes, whereas blue represents the down-regulated genes. Alt text: Heatmap of differentially expressed genes verified by qRT-PCR with color codes from red (up-regulated genes) to blue (down-regulated genes).

### Annotation and transcriptome profiling of oval and falcate conidia across different functional gene groups

To gain an overview of the encoded genes, we performed a comprehensive annotation of all predicted genes with respect to their putative roles in fungal development (conserved regulators of development, transcription factors, secondary metabolites) and pathogenicity (carbohydrate-active enzymes (CAZymes), effectors, nutrient transporters) (Figure 1b). All analyzed functional gene groups are broadly distributed across all ten major chromosomes (Chr 1–10). In contrast, the microchromosomes (MCs) in *C. graminicola* revealed that they are enriched in repeats and have reduced gene content compared to the core chromosomes. The MCs harbor a limited number of annotated genes, with approximately 84% of the total 160 genes being hypothetical. Interestingly, some of the annotated genes on the MCs are related to pathogenicity, including COGRA5_15372 (minichromosome 11), which encodes glutathionylspermidine synthase and has been previously shown to contribute to virulence in *C. graminicola* (Jaramillo et al. 2015). Our RNA-seq data analysis also revealed strong induction of COGRA5_15372 during early leaf infection (1dpi) by falcate conidia, suggesting a role in plant penetration and establishment of infection. Furthermore, the minichromosome 12 and 13 harbor 4 ankyrin repeat proteins, which are versatile protein motifs involved in mediating diverse protein-protein interactions (Bork 1993, Mosavi et al. 2002). In fungi, ankyrin repeat proteins have been shown to function as transcription factors or regulators of secondary metabolite biosynthesis, including toxins that contribute to virulence (Pedley and Walton 2001).

#### 1. Transcription factor encoding genes

Transcription factors (TFs) are proteins that bind to specific DNA sequences and regulate gene expression. All eukaryotes, including fungi, depend on appropriately orchestrated TFs to coordinate the expression of genes essential for individual developmental stages and for the induction of vital metabolic pathways. These pathways encompass carbohydrate, iron, and nitrogen metabolism, as well as tolerance to oxidative stress, osmotic conditions, pH levels and UV light. Furthermore, TFs are instrumental in establishing developmental processes like vegetative growth, tissue differentiation (fruiting bodies, conidiophores), and pathogenicity (John et al. 2021).

Analysis of the *C. graminicola* genome revealed 216 identified transcription factors (TFs) representing a diverse array of gene families (Dataset S4). The Zn_2_C_6_ (GAL4-like zinc cluster) and C_2_H_2_ zinc finger families were the most prevalent, consistent with their known abundance and regulatory significance in plant-pathogenic ascomycetes. The analysis of differentially expressed TF genes revealed that most of them exhibited higher expression in freshly harvested conidia, with a general trend towards down-regulation during subsequent development and maize leaf infection (Figure 4a). To get an overview about the most prominent gene expression changes, the 100 most variable expressed TF genes were identified using iDEP.96, highlighting several genes that showed opposite regulation patterns in oval or falcate conidia over the different analyzed stages: in freshly harvested falcate conidia, specific TFs are down-regulated, but expression increases in all developmental and pathogenicity stages. For oval conidia samples, we observe the opposite: up-regulation in freshly harvested spores and down-regulation in later stages (Figure 4b). Notable examples include the DEGs COGRA5_02342 (ascospore maturation protein; KilA-N domain), COGRA5_07325 (Zn cluster transcription factor; Zinc finger), and COGRA5_03862 (homeobox domain-containing protein; Homeobox domain). In contrast, for TFs up-regulated during oval conidia development, we did not find such clear differences to falcate conidia (Figure G File S1).

**Figure 4:**
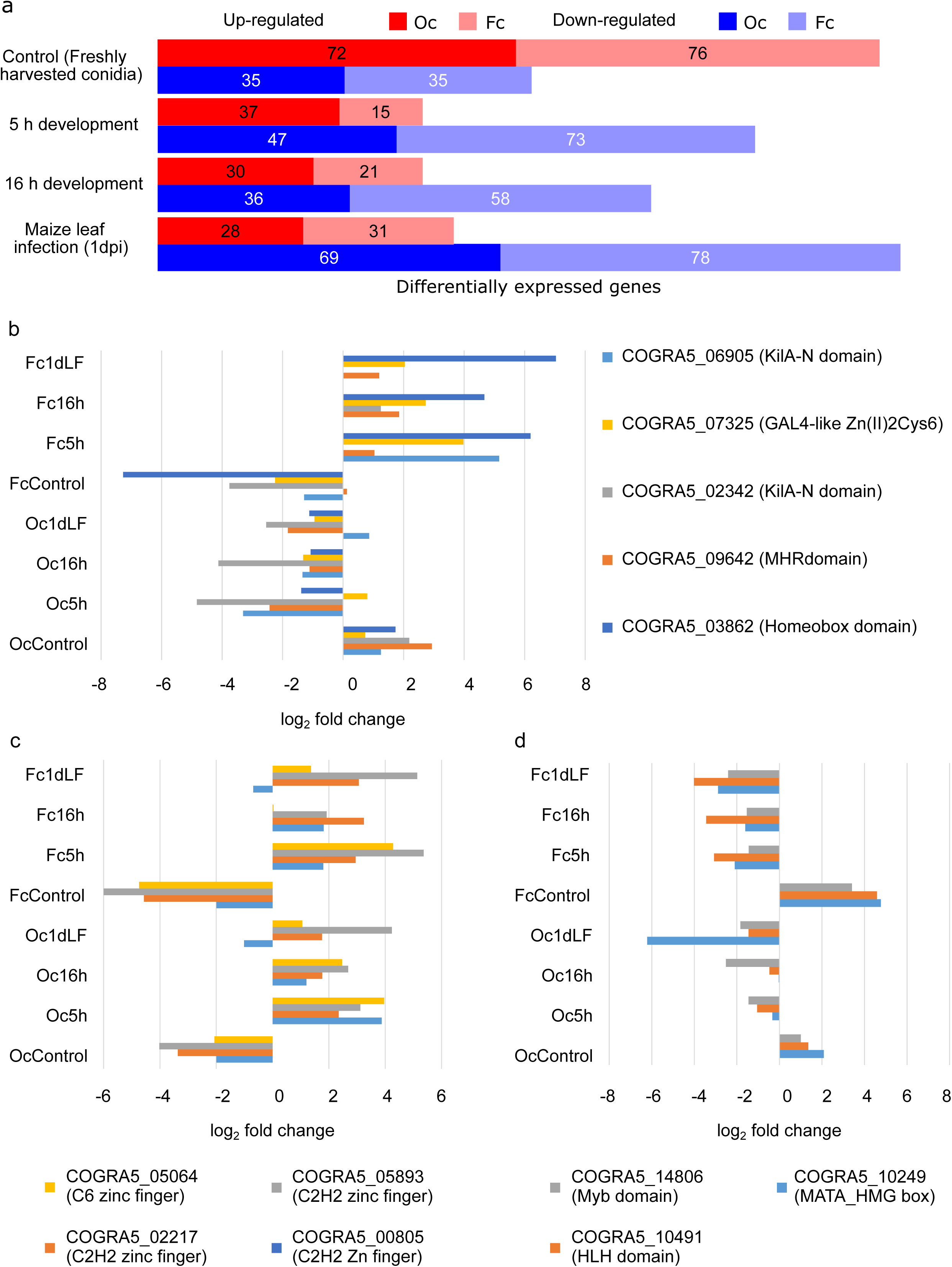
Expression patterns of transcription factor-encoding genes in oval and falcate conidia. Genes were identified using DESeq2 as differentially expressed if they had an adjusted P-value (Padj) of <0.05 and log2 values of fold expression changes were either ≥1 for up-regulated or ≤1 for down-regulated genes. As reference sample for freshly harvested conidia served mycelium sample, whereas for 5 h, 16 h, and early leaf infection, freshly harvested conidia were chosen. (a) Total number of predicted transcription factor genes showing differential regulation (log2 fold change <=-1 or >=1, padj <0.05). Red color depicts the up-regulated genes, whereas blue represents the down-regulated genes. (b-d) log2 fold change <=-1 or >=1, padj <0.05 and different color bar represent different transcription factor genes. The top 100 variable genes were identified using fold change values and were analysed using iDEP.96 (Ge et al. 2018). Hierarchical clustering was performed based on correlation distance with average linkage. Z-score normalization (centered by subtracting the mean) was applied, and a cut-off Z-score of 4 was used for analysis. (b) Transcription factor-encoding genes with induced expression for falcate conidia, as compared to oval conidia (c, d) Transcription factor genes (c) up-regulated and (d) down-regulated in both spore types during different developmental stages. Alt text: Bar charts showing differentially expressed transcription factor genes in oval and falcate conidia, with subfigures labelled from a to d, illustrating the total number of transcriptionally regulated transcription factor encoding genes and selected patterns of gene regulation.

Additionally, several TFs exhibited similar regulation patterns in both conidial types across developmental stages and leaf infection (1 dpi), such as the DEGs COGRA5_05064 (transcriptional regulatory protein Pro1, C6 zinc finger) and COGRA5_05893 (nuclear division 74 protein; C2H2 zinc finger), were consistently up-regulated during development and leaf infection (Figure 4c). Conversely, others including the DEGs COGRA5_14806 (Myb-like DNA binding protein; Myb domain) and COGRA5_10249 (female and male fertility protein; MATA_HMG box) were down-regulated in both conidia types (Figure 4d).

#### 2. Effector encoding genes

Fungal effectors are secreted proteins or small molecules that facilitate host colonization by modulating plant immune responses. These molecules are differentially expressed based on the contacting host tissue and stages of disease development in plant pathogenic species (Sonah et al. 2016). Depending on their mode of action and subcellular localization, effectors can be broadly categorized as apoplastic (extracellular) or cytoplasmic (intracellular; Todd et al. 2022).

Apoplastic effectors act in the extracellular space of plant tissues, e.g. by binding to fungal cell wall components such as chitin to prevent host recognition, whereas cytoplasmic effectors are delivered inside plant cells, where they interfere with intracellular immune signaling and defense mechanisms (Li et al. 2024). Based on the 523 annotated effector encoding genes of genome version 4 (Becerra et al. 2023), 513 putative effector genes were identified, comprising 295 apoplastic, 159 cytoplasmic and 59 apoplastic\cytoplasmic candidates (Dataset S5). To get an indication, whether effector gene expression is regulated similarly in both spore types, the Top 100 most variable genes were identified using iDEP.96. A comparison of effector gene expression patterns among those Top 100 revealed that most of those are regulated in a similar way in development and pathogenicity processes of oval and falcate conidia, reflecting that both spore types interact with the same environment and host (Figure H File S1). However, several effector genes show distinct and even opposite regulation in the 1 dpi infection stage. To analyze these initial findings in more detail, volcano plots were generated using VolcaNoseR on the basis of gene expression patterns of all effector genes (https://huygens.science.uva.nl/VolcaNoseR/). As expected, several effector genes were significantly induced during leaf infection as compared to freshly harvested conidia in both spore types (Figure I File S1). Interestingly, several of the top 10 significantly induced effector genes identified were shared among both spore types, including COGRA5_07713 (apoplastic effector), COGRA5_11054 (cytoplasmic effector), and COGRA5_03609 (apoplastic/cytoplasmic effector). However, when we used volcano plotting to visualize differences in between early leaf infection of both spore types, some effectors were visualized exhibiting opposing patterns of regulation (Figure 5, Table D File S1). For example, the known apoplastic effector COGRA5_04185 encoding a hydrophobin (Bettini et al. 2012, Eisermann et al. 2019) is strongly induced during leaf infection by falcate conidia but is not regulated in leaves infections of oval conidia. COGRA5_10559, previously described as a switch-specific effector (Kleemann et al. 2012, Oome and Van den Ackerveken 2014, Torres et al. 2016, Seidl and Van den Ackerveken 2019), was significantly up-regulated when comparing falcate and oval conidia infections.

**Figure 5:**
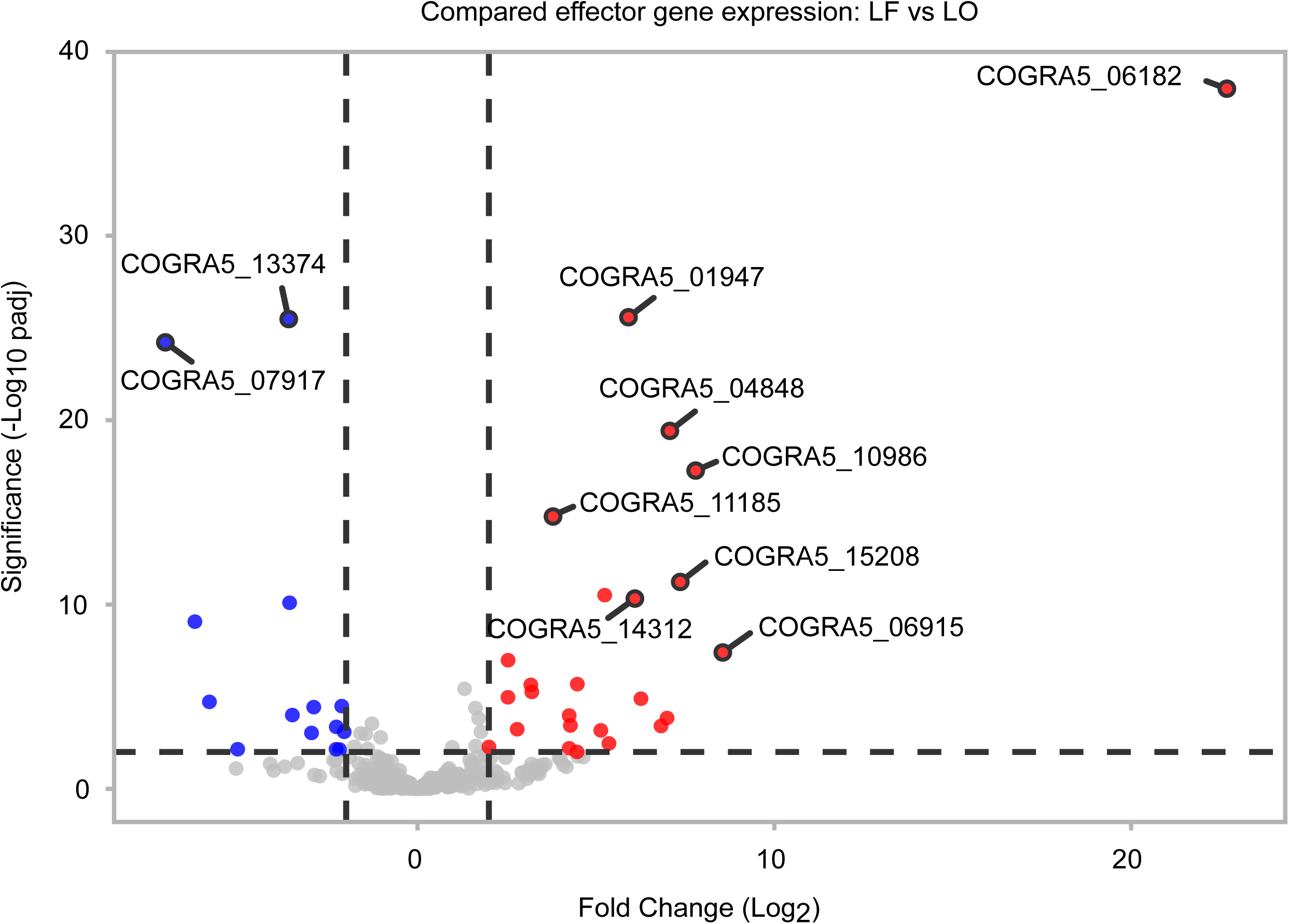
Volcano plot of effector genes during early leaf infection by oval and falcate conidia. The volcano plot was generated based on the comparison of gene expressions of falcate conidia leaf infection versus oval conidia leaf infection. VolcaNoseR settings were as follows: fold change threshold: -2 to 2; significant threshold: 2; use thresholds to annotate: changed and significant; criterion for ranking hits: Manhattan distance. Significantly regulated genes comparing falcate conidia leaf infection and oval conidia leaf infection are indicated in red (up-regulated) and blue (down-regulated). Locus tag numbers of the 10 most relevant hits are indicated. Alt text: Volcano plot showing effector gene expression differences between falcate and oval conidia during early leaf infection, with significantly regulated genes highlighted in red (up-regulated genes) and blue (down-regulated genes) and the Top 10 regulated genes indicated with locus tag numbers.

#### 3. Genes encoding membrane transporters

Transporters facilitate selective movement (import or export) of molecules across membranes. Organic and inorganic nutrients like sugars, amino acids, ions, or other metabolites, and water are likely to be taken up across the plasma membrane (Dahl et al. 2004). Transporters play a crucial role in securing survival, growth, morphogenesis, and pathogenesis by enabling the pathogen to adapt to different nutritional conditions, which are faced in and outside host cells (Verma et al. 2014, Lange and Peiter 2020). The differential expression of transporter encoding genes can therefore indicate metabolic shifts in the host, transition of fungal developmental stages, and spore type-specific physiological strategies.

From the total of 738 predicted transporter encoding genes in *C. graminicola*, including 104 newly annotated transporter genes (Dataset S6), we first analyzed the Top 100 genes showing the strongest variability in their expression, as identified by using iDEP.96. The generated heatmap depicted three main regulatory groups, which each contain about a third of all genes included in this analysis. Two groups contained transporter-encoding genes which show a general opposite regulation pattern in oval and falcate conidia over all developmental and pathogenicity stages investigated (Figure J File S1). Gene regulations in the third group do not show variability among oval and falcate conidia, but a general induction during early stages of leaf infection compared to vegetative development. Thus, these transporters thus might be involved in the adaptation of nutrition acquisition from *ex* to *in planta* environment. When we analyzed the transporter-encoding genes having the highest expression during early leaf infection in more detail, we found that within the 40 highest expressed genes there were several transporter families showing similar induction in oval and falcate conidia (Figure 6). As most prevalent, those include AAAP type amino acid permeases and sugar porters (SP). At the same time, several transporter families were over- or underrepresented when the two spore types were compared. Those include transporters from the DHA1 and DHA2 families, which are much more frequently induced in falcate than in oval conidia. Also in other respects, the profile of induced transporters differs strongly between oval and falcate conidia. 48% of the top 40 induced transporter-encoding genes in oval conidia were anion:cation symporters (ACS), whereas the same transporter family sums up to 20% in falcate conidia, indicating a probable spore type-specific strategy for the uptake of carboxylates such as succinate and citrate, organic acids and vitamins (Nie et al. 2017). We further identified several other transporter gene families that show clear spore type-specific expression patterns, such as monocarboxylate porter (MCP), p-ATPases, and tricarboxylate carriers, which are prominently up-regulated in falcate conidia, indicating adaptation to a nutrient environment rich in carboxylic acids (like C4 leaves). In oval conidia, several members from the oligopeptide transporter (OPT) and L-type amino acid transporter (LAT) families were induced. Since these were absent among induced transporter genes in falcate conidia, our results point to a preference of these nitrogen sources in oval conidia.

**Figure 6:**
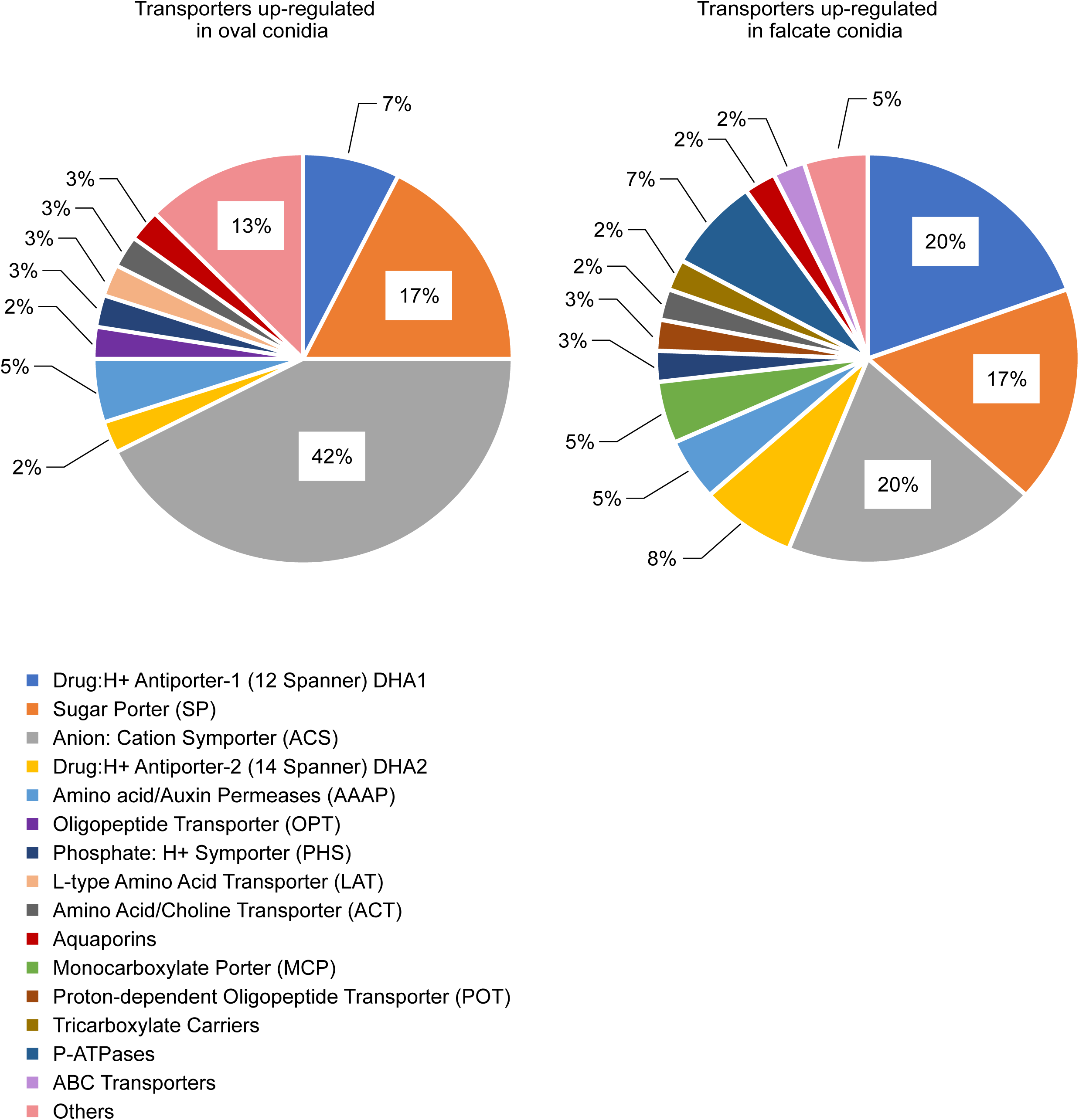
Differentially expressed transporter genes showing induced expression during early leaf infection. Genes were identified as differentially expressed if they had an adjusted P-value (Padj) of <0.05 and log2 values of fold expression changes were either ≥1 for up-regulated or ≤1 for down-regulated genes. Transporter gene families with elevated expression in falcate conidia and oval conidia, identified among the top 40 expressed genes during early leaf infections. Alt text: Pie charts showing transporter gene families with significantly induced expression during early leaf infection in oval and falcate conidia.

#### 4. Genes encoding CAZymes

Microorganisms produce a diverse repertoire of carbohydrate-active enzymes (CAZymes) involved in the synthesis, modification, and degradation of complex carbohydrates (Wardman and Withers 2024). These enzymes hydrolyze major plant cell wall polymers such as cellulose, hemicellulose, and pectin, enabling the utilization of plant biomass as a nutrient source. CAZymes are organized into distinct families based on protein sequence similarity and conserved three-dimensional structural folds (Barrett et al. 2020b).

In total, 663 CAZymes were annotated in the new genome version 5, out of which 558 are differentially expressed in at least one pairwise comparison (O_vs_M, F_vs_M, LO_vs_O, O5_vs_O, O16_vs_O, LF_vs_F, F5_vs_F, F16_vs_F, LF_vs_LO; Dataset S7). This is an increase from the previously annotated 467 CAZymes, highlighting the complexity and diversity of CAZyme regulation in this fungus (Baroncelli et al. 2016). To reveal CAZyme families showing a high level of regulation during development and pathogenicity, the Top 100 variable genes were analyzed in detail. In general, similar patterns to the evaluation of transporter-encoding genes can be observed: besides CAZmyes regulated in an opposite way in oval and falcate conidia, there are several showing a general upregulation during early leaf infection stage in both spore types (Figure 7 a, b; Figure K File S1). The latter group thus might stand for a general adaptation of the fungus from saprophytic to pathogenic growth, in which predominantly complex substrates from plants are used for nutrition. Among the families showing strict spore type specificity there were, for example, Carbohydrate Esterase family CE16 (acetylesterase) uniquely up-regulated in falcate conidia, whereas Auxiliary Activity famiy AA12+AA8 (oxidoreductase) was specifically induced in oval conidia. The most striking distinction between spore types was observed in the Auxiliary Activities (AA) family (Figure 7, Figure L File S1). In falcate conidia, members of the AA9 family (lytic polysaccharide monooxygenases with cellulase and xylanase activities) were strongly up-regulated, alongside higher expression of Glycoside Hydrolase family GH10 cellulase/xylanase genes (Figure L of File S1). These results suggest that falcate spores may be primed for the rapid breakdown of cellulose- and hemicellulose-rich plant cell walls, facilitating host tissue penetration and enabling nutrient release during the early necrotrophic phase of infection. In contrast, oval conidia exhibited elevated expression of Polysaccharide Lyase family PL3 (pectate lyase) during early developmental stages (5 h and 16 h; Figure M of File S1). The indicated specialization for the degradation of pectin, an abundant component of the middle lamella, could support more targeted and localized modification of host cell walls by oval conidia, potentially aiding in adhesion or subtle wall loosening.

**Figure 7:**
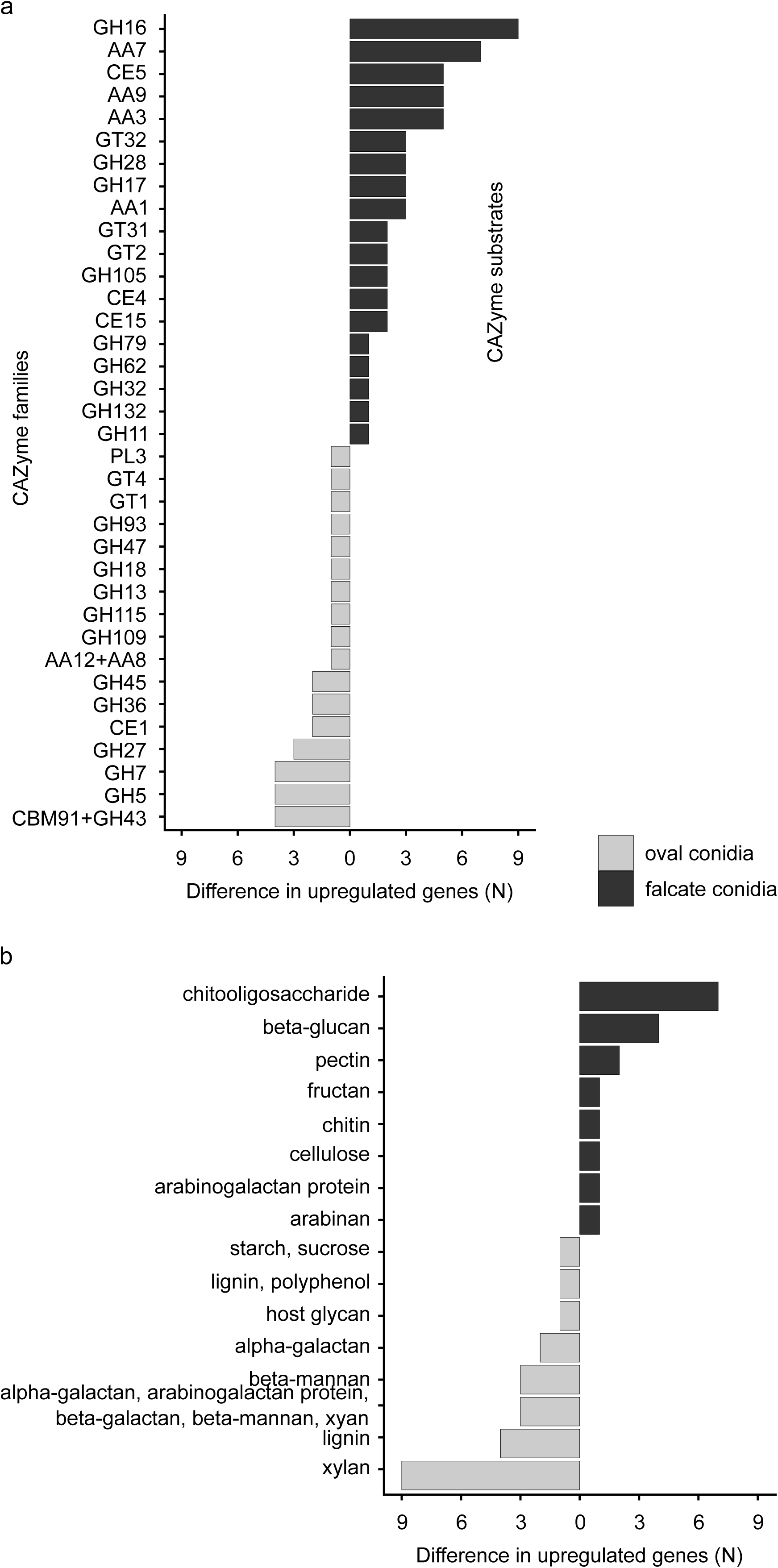
Most variable differentially expressed Carbohydrate-Active Enzymes (CAZymes) in *C. graminicola* asexual spore types during leaf infection. As reference condition served freshly harvested conidia of the corresponding spore type. (a) Distribution of CAZyme families and (b) corresponding predicted CAZyme substrate categories associated, up-regulated in oval and falcate conidia. Identification of the top 100 variable genes was performed using iDEP.96 on the basis of differentially expressed genes identified (DEseq). Hierarchical clustering was performed based on correlation distance with average linkage. Genes were identified as differentially expressed if they had an adjusted P-value (Padj) of <0.05 and log2 values of fold expression changes were either ≥1 for up-regulated or ≤1 for down-regulated genes. Alt text: Expression patterns of the 100 most variable CAZyme genes in *C. graminicola* leaf infection with oval and falcate conidia, with subfigures labelled from a to b, displaying CAZmye families and the corresponding substrates.

#### 5. Genes encoding regulators of development

In fungi, the regulatory proteins coordinating crucial processes such as growth, spore formation (conidiogenesis), morphological differentiation, reproduction, and development of specialized structures like fruiting bodies and plant surface penetrating structures, are highly conserved (Nolting and Pöggeler 2006, Wong Sak Hoi and Dumas 2010, Lichius and Lord 2014, Turrà et al. 2014). For this study, we verified the presence of 122 conserved regulators of development using BLAST analysis based on *Sordaria macrospora and Aspergillus flavus* NRRL3357 protein sequences (Dataset S8; Teichert et al. 2020, Cho et al. 2022). Most developmental genes did not show a high level of regulation at the different timepoints investigated. Genes which showed a regulation pattern, however, exhibited a general downregulation in freshly harvested spores and an induction of expression during the progression of vegetative development and pathogenicity.

Among the major regulated genes, the ones encoding enzymes in the melanin biosynthesis were up-regulated in both spore types during leaf infection (Figure N File S1). This aligns with previous findings that melanin functions are an important virulence factor in plant-pathogenic fungi, facilitating early host penetration, stress adaptation, and evasion of plant defenses (Howard and Valent 1996, Li et al. 2024). Similarly, we observed induction of Rho4-like GTPase, which plays a crucial role in various cellular processes, including β-1,3-glucan synthesis, cell wall integrity, growth of vegetative hyphae, conidiation, infection structure differentiation, and is also required for full virulence, in both spore types during early leaf infections (Oliveira-Garcia et al. 2022). For fungal NADPH oxidase complexes (NOX), we found very distinct expression patterns: whereas subunits required for the assembly of the NOX1 complex (catalytic subunit Nox1 (COGRA5_02814), NoxD regulator (COGRA5_05373) showed a strong regulation depending on the developmental stage or spore type, the catalytic subunit of the NOX2 complex, Nox2 (COGRA5_09929), showed hardly any gene regulation. This is interesting, since both complexes are associated with very different developmental processes in fungi, albeit both are generating signaling competent reactive oxygen species (Siegmund et al. 2015, Li et al. 2022): NOX1 was associated in with successful germling fusion, vegetative growth and sexual development, whereas NOX2 is required for ascospore germination and chemotropic sensing of root derived class III peroxidases in several fungi. In interactions with the host, however, both NOX complexes serve the penetration of the plant surface (Cano-Domínguez et al. 2008, Kayano et al. 2013, Ryder et al. 2013, Dirschnabel et al. 2014, Marschall et al. 2016, Nordzieke et al. 2019a, Rudolph et al. 2024). Thus, one might expect differential patterns of *Cgnox1* and *Cgnox2* gene expressions in the different samples investigated. The absence of a clear transcript regulation for *Cgnox2* might thus indicate that the activity of the NOX2 complex is not regulated primarily by gene expression changes, but by the assembly of the complex components.

#### 6. Secondary metabolite biosynthesis genes

Secondary metabolites (SMs) are low molecular weight organic compounds produced by fungi, plants and bacteria. In contrast to primary metabolites essential for the organisms’ survival, secondary metabolites are produced in specific environmental and developmental conditions to improve ecological fitness (Bills and Gloer 2016, Macheleidt et al. 2016). Primary metabolites originating from central metabolic pathways serve as essential building blocks together with Acyl-CoAs and amino acids (Macheleidt et al. 2016, Keller 2019). Fungal SMs are chemically diverse and fall into four major classes: polyketides, terpenoids, phenolics, and non-ribosomal peptides (Pusztahelyi et al. 2015). Genes responsible for SM biosynthesis are typically organized in contiguous order as biosynthetic gene clusters (BGC; Keller 2019).

Regulation of BGCs is closely related to the ecological context and infection stage, underlining the adaptive significance of secondary metabolism during pathogenesis (Lysøe et al. 2011, Keller 2019).

Based on the *C. graminicola* genome sequence V5, gene candidates belonging to SM biosynthesis clusters were predicted using antiSMASH version fungiSMASH 7.0.0 (Blin et al. 2021). Among these, 466 genes are differentially regulated during various developmental stages and early stages of maize leaf infection 1 dpi (Dataset S9). Notably, 193 SM BGCs displayed a decreased expression in freshly harvested samples of both falcate and oval conidia. Conversely, 133 BGCs exhibited induction during leaf infection in both spore types (Figure O File S1). Notably, the fusaridione A cluster, which was originally identified in *Fusarium heterosporumin,* was induced upon host interaction, aligning with previous reports (Kakule et al. 2013). A similar pattern is displayed by the depudecin BGC. This cluster enables biosynthesis of the histone deacetylase inhibitor depudecin, which contributes to virulence in *Alternaria brassicicola* (Wight et al. 2009, Pedras and Park 2015). Additionally, the BGC of three antimitotic mycotoxins, phomopsins A, B, and E display increased expression during host interaction, similarly to observations from *Phomopsis leptostromiformis* (Hamel 1992, Grimley et al. 2007, Izuchi et al. 2011).

## Discussion

In this study, we performed a comprehensive RNA-seq analysis based on genome resequencing of our laboratory strain of the corn anthracnose fungus *C. graminicola* which is derived from the original isolate CgM2/M1.001. We focused on the elucidation of spore type-specific gene expression patterns of critical stages in development and early leaf infection of oval and falcate conidia and their derived hyphae. For data interpretation, we concentrated on six functional gene groups, comprising genes relevant as transcription factors, effectors, transporter proteins, CAZymes, developmental regulators, and as parts of secondary metabolite gene clusters. Overall, we found gene expression changes of both conidia types regarding germination and germling stages, along with a general increase in gene expression levels observed during early leaf infections. In comparison to mycelium samples, freshly harvested oval and falcate conidia showed clear differences in gene expression patterns, with overlaps between the two conidia types but also spore type-specific gene expression changes are evident (Figure 2). Similarly, we found that during early maize leaf infection the top variable genes encoding effector proteins, transporters, and CAZymes show uniform expression patterns, reflecting the general adaptation to a host environment. At the same time, several genes are specifically regulated in oval and falcate conidia, some with even opposite expression patterns (up-regulation in oval conidia and down-regulation in falcate conidia and vice versa), pointing to probable spore type-specific processes during host interactions (Figures H, J, and K File S1). Together, these findings indicate that oval and falcate-shaped spores express a core gene set responsible for their identity as conidia, clearly different from vegetative mycelium. On the other hand, both conidia types have an individual signature, which is maintained also during early maize leaf infection. These findings support earlier findings from phenotypic investigations, in which oval and falcate conidia were shown to undergo specific developmental processes and explore unique infection strategies meanwhile interacting with host leaves (Nordzieke et al. 2019b). Altogether, this study provides a comprehensive overview about gene expression changes in vegetative development and during early leaf interaction of oval and falcate conidia, forming the basis for spore type-specific development and virulence of *C. graminicola*.

In previous studies, genome sequencing in combination with RNA-seq was used to understand critical factors determining the symptoms developed by closely related plant pathogens. For example, in *Zymoseptoria*, a genus of plant pathogenic fungi specialized in the infection of grasses, comparison of the genomes of four different species identified species-specific orphan genes and genes encoding small, secreted proteins with putative functions for virulence and host specificity (Grandaubert et al. 2015). Furthermore, RNA-seq analysis of eight *Fusarium fujikuroi* isolates revealed a correlation between the formation of secondary metabolites and two distinct pathotypes on rice. In this case, gibberellic acid (GA) biosynthesis is positively correlated with typical symptom development of *bakanae* disease like hyper-elongated seedlings, while fumosin-generating strains showed stunting and early withering of infected seedlings (Niehaus et al. 2017). Similarly, RNA-seq can give important hints to the regulation of plant defense responses during pathogenic interactions. For example, comparison of gene expression profiles of soybeans infected with non-pathogenic and pathogenic strains of *Fusarium oxysporum* revealed high numbers and magnitudes of differentially expressed genes (HDEGs) in the plant comparing both infection scenarios (Lanubile et al. 2015). In all those examples, different isolates of the same pathogenic species or closely related species were analyzed, having isolate-specific or species-specific genetic information encoded in their genomes, respectively. Whereas comparable studies are numerous, distinct transcriptomes of asexual spore types of a single species with the same genetic information are underexplored. In one case, the transcriptomes of freshly harvested microconidia and macroconidia of the rice blast fungus *Magnaporthe oryzae* were analyzed, providing evidence that only seven genes are specifically expressed in microconidia and eight additional genes show increased expression in macroconidia after harvest (Zhang et al. 2014). In our study, the numbers of differentially regulated genes identified in freshly harvested oval and falcate conidia of *C. graminicola* were much higher (Figure 2): compared to mycelium samples, oval and falcate conidia showed increased expression of 1483 genes in common and, additionally, similar numbers of up-regulated spore type-specific genes (oval conidia: 1405, falcate conidia: 1799). We observed a similar pattern when comparing down-regulated genes respective to mycelium samples: in both conidia types, 1791 genes are down-regulated, but we observed >1000 genes specifically for oval (1189) and falcate conidia (1411). Bearing in mind that the genome of *C. graminicola* encodes 15,481 genes, we observed roughly 30% of the genes up-regulated and 28% of genes down-regulated in freshly harvested oval and falcate conidia, underpinning the metabolic activity of both spore types. Monitoring later stages of development, we observed rapid transcriptional changes in both spore types after 5 h, although only oval conidia showed germination under the conditions monitored and falcate conidia remained dormant. These surprising observations, however, are in line with a previous report regarding transcriptional activity of dormant *Aspergillus* conidia, providing evidence that fungal conidia are able to react to environmental signals, shaping the transcripts and thus the later behavior after dormancy breaking (Wang et al. 2021).

Apart from general gene expression changes, we identified numerous genes with similar, specific or even opposite regulations in oval and falcate conidia, of which we further analyzed a subset using qRT-PCR (Figure 3). For instance, COGRA5_11040, encoding an Aspzincin_M35 metallopeptidase, is highly induced in both oval and falcate conidia during early leaf infection (1 dpi). Since metalloproteinases degrade proteins in host tissues and increase thereby the hosts’ susceptibility to disease, this result might indicate that oval and falcate conidia use the same strategy for masking their presence (Huang et al. 2020, Wang et al. 2024). In contrast, Lysine-domain containing proteins (LysMs) were mainly induced during leaf infection with falcate conidia, but to a lesser extent with oval conidia (Dataset S10). However, the LysM encoding gene COGRA5_04487 even shows an opposite regulation. Fungal LysM proteins are effectors and play crucial roles in regulating development, maintaining cell wall integrity by binding chitin, and contributing to pathogenicity by suppressing host defenses and facilitating host colonization (Muraosa et al. 2019, Dubey et al. 2020). In contrast to those findings, most of the *TauD* genes present in *C. graminicola* genome are highly expressed during oval conidia leaf infection, but not in falcate conidia (Figure 3; Dataset S3). TauD proteins enable fungi to survive in nutrient-deprived conditions by utilizing taurine and xanthine as nutrient sources (Eichhorn et al. 1997, Montero-Morán et al. 2007). This is consistent with our previous results that oval conidia are able to survive and germinate in nutrient-deprived conditions, but falcate conidia are not (Nordzieke et al. 2019b, Rudolph et al. 2025b). Intriguingly, nutrient starvation induced by cultivation media or the inoculation on intact plant surfaces induce fusion of oval conidia derived germlings, which give rise to hyphopodia formation for plant penetration (Nordzieke et al. 2019b). *TauD* genes therefore could contribute to successful germination and germling fusion during colony formation and the pathogenicity of *C. graminicola*.

Additionally, our RNA-seq analysis indicated that the two spore types of *C. graminicola* use distinct gene repertoires to adapt to the shift from saprotrophic growth to pathogenic interaction with the plant. For example, CAZymes and transporters, which are required for nutrient acquisition and uptake, comprise family members with a specific upregulation during leaf infections in both spore types (Figures J and K File S1). At the same time, there are several CAZyme and transporter-encoding genes, which are oppositely regulated in oval and falcate conidia, meaning that gene up-regulation in oval conidia samples parallels down-regulation in falcate conidia. Such regulation indicates that these genes might be vital for nutrition of a distinct spore type. For example, the CAZyme families AA9 and GH10 are induced in falcate conidia during early leaf infections. Together, these enzymes enable rapid breakdown of cellulose- and hemicellulose rich plant cell walls and facilitate tissue penetration and nutrient release in early infection (Bradley et al. 2022; Figure S10). At the same time, falcate conidia show an elevated number of DHA transporters among the top regulated genes during early leaf infections, indicating an increased capacity for the detoxification of host defense metabolites (Cavalheiro et al. 2018). In contrast, oval conidia express oligopeptide transporters (OPT) and L-type amino acid transporters (LAT), which are potentially involved in organic nitrogen uptake from the host (Reuß and Morschhäuser 2006, Kantipudi and Fotiadis 2021; Figure 6). They also showed elevated expression of PL3 (pectate lyase) during early vegetative development (5 h and 16 h), which degrades pectin in the middle lamella of plant cells, supporting adhesion, controlled wall loosening, and early biotrophic establishment without extensive damage (Figure 7, Figure L File S1; Yang et al. 2018). Focusing on expressed effector genes, we observed the same general pattern: several effector encoding genes were induced upon the initiation of host leaf infections in oval and falcate conidia (Figures H and I Files S1), indicating a general role in the interaction with the host. Despite those, there are some effector genes, which are differentially regulated when we compare gene expressions between early leaf infection of falcate and oval conidia (Figure 5, Table D File S1). One of those, COGRA5_10559, was previously characterized as necrotrophy-associated, switch-specific effector in *C. graminicola* falcate conidia (Kleemann et al. 2012, Oome and Van den Ackerveken 2014, Torres et al. 2016, Seidl and Van den Ackerveken 2019). Interpreting these data, it is however important to keep in mind the different strategies for leaf infection explored by both conidia types: oval conidia form germling networks on leaves, whereas falcate conidia form appressoria from short germ tubes. Independent on the infection process, both spore types are therefore in different physiological stages, which might explain some of the transcriptomal changes seen during early leaf infections. Future studies will have to reveal whether the conidia-specific gene expression patterns seen have consequences for the maize infection processes induced by *C. graminicola* oval and falcate conidia.

## Acknowledgements

We thank Gabriele Beyer and Gertrud Stahlhut for excellent technical assistance and Simone Lewandowski for support with some of the experiments. This work was funded by the Deutsche Forschungsgemeinschaft (Bonn-Bad Godesberg). Grants were provided to D.E.N. (IGRK2172 “PRoTECT” – Projektnummer 273134146), to M.A.G. (Grant No. GU 2252/1-1, Project No. 460261834), and to M.N. (project number NO407/8-1). Additional funding was from a discovery grant from the Natural Sciences and Engineering Research Council of Canada (NSERC) (to J.K.) and by the NSERC-CREATE Program contribution to the PRoTECT program. J.K. is the Power Corporation Fellow of the Canadian Institute for Advanced Research (CIFAR) program on the Fungal Kingdom: Threats & Opportunities. We further thank the Max Planck Institute for Evolutionary Biology for the computing infrastructure. D. R. was supported by the GGNB program PRoTECT. The manuscript text was edited to improve spelling and grammar using the AI tools ‘Chat AI’ (version v0.9.0) of, Gesellschaft für wissenschaftliche Datenverarbeitung mbH Göttingen‘, GWDG and ‘Academic AI’ of ACOnet, Austria.

## Author contributions

DR provided raw material for genome sequencing and to generate RNA-seq data, performed data interpretation, verified the results via qRT-PCR and wrote the manuscript. AP performed genome resequencing, quality checks and sequence trimming of Illumine and Nanopore reads. MN performed genome assembly and bioinformatical RNA-seq data analysis. JK and MK assisted in performing qRT-PCR and data analysis. Annotation of functional gene groups was conducted by SP, LV, MG, KA, DR and DN. Specifically, SP was involved in annotation of development regulators, LV in membrane transporters, MG in CAZymes, KA in secondary metabolite gene clusters, DN in effector proteins and DR in transcription factors and developmental genes. AP, RD, SP, LV, MG, KA, JK, MK, DR and DN reviewed the manuscript. DN designed and supervised the project, contributed in data interpretation, edited the manuscript, and was responsible for project funding.

## Data availability

The data underlying this article are available in the European Nucleotide Archive (ENA; https://www.ebi.ac.uk/ena/browser/home) and can be accessed under the project numbers PRJEB101887 (genome resequencing), PRJEB104144 (RNA-seq), and ERZ29778039 (genome annotation). The annotated genome files including genome assembly, annotation, and sequences for each contig have been added as supplementary information (file S3.zip). For submission of supplementary datasets, main Figures, File S3.zip and graphical abstract, GSA Figshare portal was used. That submission contains the following datasets: Dataset S1: Annotation of protein coding genes of *Colletotrichum graminicola* M1.001 (CgM2 wildtype strain) with new gene IDs (COGRA5); Dataset S2: RNA-seq data of all the *Colletotrichum graminicola* M1.001 (CgM2 wildtype strain) genes labelled with different locus tags; Dataset S3: Expression of Taurine catabolism dioxygenase (TauD) genes in *Colletotrichum graminicola* M1.001 (CgM2 wildtype strain); Dataset S4: RNA-seq data of predicted transcription factors in *Colletotrichum graminicola* M1.001 (CgM2 wildtype strain), identified using data from 312 predicted transcription factor genes in *Neurospora crassa* OR74A (Carrillo et al. 2017); Dataset S5: RNA-seq data of predicted effector genes in *Colletotrichum graminicola* M1.001 (CgM2 wildtype strain); Dataset S6: RNA-seq data of predicted membrane transport proteins in *Colletotrichum graminicola* M1.001 (CgM2 wildtype strain); Dataset S7: RNA-seq data of predicted CAZymes in *Colletotrichum graminicola* M1.001 (CgM2 wildtype strain); Dataset S8: RNA-seq data of predicted developmental genes in *Colletotrichum graminicola* M1.001 (CgM2 wildtype strain), identified using data based on *Sordaria macrospora* and *Aspergillus flavus* NRRL3357 protein sequences (Teichert et al. 2020, Cho et al. 2022); Dataset S9: RNA-seq data of predicted secondary metabolites biosynthesis clusters in *Colletotrichum graminicola* M1.001 (CgM2 wildtype strain), which was predicted using antiSMASH version fungiSMASH 7.0.0 (Blin et al. 2021). Dataset S10: Expression of LysM candidates (CBM 50) in *Colletotrichum graminicola* M1.001 (CgM2 wildtype strain); Figure 1: Overview of the *C. graminicola* assembly V5; Figure 2: Differentially expressed genes in two different conidia types identified by DESeq2; Figure 3: Comparative analysis of transcriptomic expression using both qRT-PCR and RNA-seq of selected genes; Figure 4: Expression patterns of transcription factor-encoding genes in oval and falcate conidia; Figure 5: Volcano plot of effector genes during early leaf infection by oval and falcate conidia; Figure 6: Differentially expressed transporter genes showing induced expression during early leaf infection; Figure 7: Most variable differentially expressed Carbohydrate-Active Enzymes (CAZymes) in *C. graminicola* asexual spore types during leaf infection; File S3.zip: Annotated genome files including genome assembly, annotation, and sequences for each contig have been added as supplementary information; graphical abstract: graphical abstract;

## Supplementary Dataset Headers

Dataset S1: Annotation of protein coding genes of *Colletotrichum graminicola* M1.001 (CgM2 wildtype strain) with new gene IDs (COGRA5).

Dataset S2: RNA-seq data of all the *Colletotrichum graminicola* M1.001 (CgM2 wildtype strain) genes lablled with different locus tags.

Dataset S3: Expression of Taurine catabolism dioxygenase (TauD) genes in *Colletotrichum graminicola* M1.001 (CgM2 wildtype strain).

Dataset S4: RNA-seq data of predicted transcription factors in *Colletotrichum graminicola* M1.001 (CgM2 wildtype strain), identified using data from 312 predicted transcription factor genes in *Neurospora crassa OR74A* (Carrillo et al. 2017).

Dataset S5: RNA-seq data of predicted effector genes in *Colletotrichum graminicola* M1.001 (CgM2 wildtype strain).

Dataset S6: RNA-seq data of predicted membrane transport proteins in *Colletotrichum graminicola* M1.001 (CgM2 wildtype strain).

Dataset S7: RNA-seq data of predicted CAZymes in *Colletotrichum graminicola* M1.001 (CgM2 wildtype strain).

Dataset S8: Expression of LysM candidates (CBM 50) in *Colletotrichum graminicola* M1.001 (CgM2 wildtype strain).

Dataset S9: RNA-seq data of predicted developmental genes in *Colletotrichum graminicola* M1.001 (CgM2 wildtype strain), identified using data based on *Sordaria macrospora* and *Aspergillus flavus* NRRL3357 protein sequences (Teichert et al. 2020, Cho et al. 2022).

Dataset S10: RNA-seq data of predicted secondary metabolites biosynthesis clusters in *Colletotrichum graminicola* M1.001 (CgM2 wildtype strain), which was predicted using antiSMASH version fungiSMASH 7.0.0 (Blin et al. 2021).

## Supplementary Figure legends (File S1)

Figure A: Overview of RNA samples prepared. From each sample, including *C. graminicola* mycelium (a), falcate conidia (b), and oval conidia (c), 3 replicates were prepared. (b, c) Conidia samples were taken after different incubation timepoints including freshly harvested conidia, conidia incubated for the indicated timepoints under nutrient limiting conditions (23°C), and early maize leaf infection stages (1 dpi).

Alt text: Sample overview, with subfigures labelled from a to c, displaying the different samples used for the transcriptome analysis.

Figure B: Heatmap displaying the expression profiles of the 100 most variable genes in freshly harvested falcate and oval conidia. Red indicates up-regulation and blue indicates down-regulation of gene expression. The top 100 variable genes were identified using fold change values and were analyzed using iDEP.96 (Ge et al. 2018) where log2 fold change <=-1 or >=1, padj <0.05 was used. Hierarchical clustering was performed based on correlation distance with average linkage. Z-score normalization (centered by subtracting the mean) was applied, and a cut-off Z-score of 4 was used for analysis.

Alt text: Heatmap showing the expression profiles of the most regulated genes (Top 100) in falcate and oval conidia after harvest, with color codes from red (up-regulated genes) to blue (down-regulated genes).

Figure C: 100 most variable genes in falcate and oval conidia following 5 h of incubation at nutrient limiting conditions depicted in a heatmap. Expression levels are color-coded, with red representing up-regulated and blue representing down-regulated genes. The top 100 variable genes were identified using fold change values and were analyzed using iDEP.96 (Ge et al. 2018) where log2 fold change <=-1 or >=1, padj <0.05 was used. Hierarchical clustering was performed based on correlation distance with average linkage. Z-score normalization (centered by subtracting the mean) was applied, and a cut-off Z-score of 4 was used for analysis.

Alt text: Heatmap showing the expression profiles of the most regulated genes (Top 100) in falcate and oval conidia after 5 h of cultivation in nutrient-limiting conditions, with color codes from red (up-regulated genes) to blue (down-regulated genes).

Figure D: Heatmap displaying 100 most variable genes in falcate and oval conidia following 16 h of incubation at nutrient limiting conditions. Up-regulated genes are depicted in red and down-regulated in blue. The top 100 variable genes were identified using fold change values and were analyzed using iDEP.96 (Ge et al. 2018) where log2 fold change <=-1 or >=1, padj <0.05 was used. Hierarchical clustering was performed based on correlation distance with average linkage. Z-score normalization (centered by subtracting the mean) was applied, and a cut-off Z-score of 4 was used for analysis.

Alt text: Heatmap showing the expression profiles of the most regulated genes (Top 100) in falcate and oval conidia after 16 h of cultivation in nutrient-limiting conditions, with color codes from red (up-regulated genes) to blue (down-regulated genes).

Figure E: Heatmap displaying 100 most variable genes in falcate and oval conidia harvested from maize leaves after 1 d of inoculation. The heatmap uses red and blue to denote up-regulation and down-regulation, respectively. The top 100 variable genes were identified using fold change values and were analyzed using iDEP.96 (Ge et al. 2018) where log2 fold change <=-1 or >=1, padj <0.05 was used. Hierarchical clustering was performed based on correlation distance with average linkage. Z-score normalization (centered by subtracting the mean) was applied, and a cut-off Z-score of 4 was used for analysis.

Alt text: Heatmap showing the expression profiles of the most regulated genes (Top 100) in falcate and oval conidia after 1 d of leaf inoculation, with color codes from red (up-regulated genes) to blue (down-regulated genes).

Figure F: Expression analysis of selected differentially expressed genes with qRT-PCR. RNA was taken from the same samples which were analyzed in the RNA-seq analysis. After DNA digestion, reverse transcription was used to generate cDNA, serving as template for the qRT-PCR reaction. Correct performance of the used oligonucleotides was tested beforehand. Base means of the corresponding genes were: 388.88 (COGRA5_13220), 1152.99 (COGRA5_00514), 121.67 (COGRA5_09579), 260.96 (COGRA5_06545), 371.98 (COGRA5_00442), 227.90 (COGRA5_11040), 145.46 (COGRA5_14555).

Alt text: Bar chart showing qRT-PCR expression levels for selected differentially expressed genes.

Figure G: Differentially expressed transcription factor-encoding genes showing increased expression in oval conidia relative to falcate conidia compared to the corresponding refence conditions across different developmental stages. As reference conditions for freshly harvested conidia (FcControl, OcControl) served mycelium, for all developmental and pathogenicity-related conditions (Fc5h, Fc16h, Fc1dLF, Oc5h, Oc16h, Oc1dLF) the corresponding freshly harvested conidia samples were chosen. The top 100 variable genes were identified using fold change values and were analyzed using iDEP.96 (Ge et al. 2018) where log2 fold change <=-1 or >=1, padj <0.05 was used. Hierarchical clustering was performed based on correlation distance with average linkage. Z-score normalization (centered by subtracting the mean) was applied, and a cut-off Z-score of 4 was used for analysis. Here different color bars represent different transcription factor genes.

Alt text: Bar chart showing differentially expressed transcription factor-encoding genes with higher expression in oval conidia than in falcate conidia.

Figure H: Heatmap to illustrate the 100 most variable expressed effector genes at different developmental stages in two conidia types. Heatmap was created using the iDEP.96 based on the corresponding log2 fold change values. Red color depicts the up-regulated genes, whereas blue represents the down-regulated genes. Hierarchical clustering was performed based on correlation distance with average linkage. Z-score normalization (centered by subtracting the mean) was applied, and a cut-off Z-score of 4 was used for visualization.

Alt text: Heatmap showing the expression profiles of the most regulated effector-encoding genes (Top 100) in falcate and oval conidia, with color codes from red (up-regulated genes) to blue (down-regulated genes).

Figure I: Regulation of effector-encoding genes during early leaf infection. The volcano plots were generated based on the comparison of gene expression data of falcate conidia leaf infection vs freshly harvested falcate conidia (a) and oval conidia leaf infection vs freshly harvested oval conidia (b). VolcaNoseR settings were as followed: fold change threshold: -2 to 2; significant threshold: 2; use thresholds to annotate: changed and significant; criterion for ranking hits: Manhattan distance. Significantly regulated are indicated in red (up-regulated) and blue (down-regulated). Locus tag numbers of the 10 most relevant hits are indicated.

Alt text: Volcano plots showing regulation of effector-encoding genes during early maize leaf infection in oval and falcate conidia, with subfigures labelled from a to b, illustrating expression changes during early leaf infection with freshly harvested conidia as reference samples.

Figure J: Heatmap to illustrate the 100 most variable transporter-encoding genes at different developmental stages in two conidia types. Heatmap was created using the iDEP.96 based on the corresponding log2 fold change values. Red color depicts the up-regulated genes, whereas blue represents the down-regulated genes. Hierarchical clustering was performed based on correlation distance with average linkage. Z-score normalization (centered by subtracting the mean) was applied, and a cut-off Z-score of 4 was used for visualization.

Alt text: Heatmap showing the expression profiles of the most regulated transporter-encoding genes (Top 100) in falcate and oval conidia, with color codes from red (up-regulated genes) to blue (down-regulated genes).

Figure K: Heatmap to illustrate the 100 most variable genes encoding for Carbohydrate-Active Enzymes (CAZymes) at different developmental stages in two conidia types. Heatmap was created using the iDEP.96 based on the corresponding log2 fold change values. Red color depicts the up-regulated genes, whereas blue represents the down-regulated genes. Hierarchical clustering was performed based on correlation distance with average linkage. Z-score normalization (centered by subtracting the mean) was applied, and a cut-off Z-score of 4 was used for visualization.

Alt text: Heatmap showing the expression profiles of the most regulated CAZyme-encoding genes (Top 100) in falcate and oval conidia, with color codes from red (up-regulated genes) to blue (down-regulated genes).

Figure L: Most variable differentially expressed Carbohydrate-Active Enzymes (CAZymes) in *C. graminicola* asexual spore types during leaf infection. Expression dynamics (log₂ fold change) of major CAZyme classes across different developmental stages and early maize leaf infection (1 dpi). As reference conditions served freshly harvested oval and falcate conidia for all time points tested for the corresponding spore type. Statistical significance was tested using a Wilcoxon test, p-values were adjusted with the Holm method. Significances displayed give differences between conidial types within each CAZyme class at the corresponding developmental stage, n = numbers of genes included. Identification of the top 100 variable genes was performed using iDEP.96 based on differentially expressed genes identified (DEseq). Hierarchical clustering was performed based on correlation distance with average linkage. Z-score normalization (centered by subtracting the mean) was applied, and a cut-off Z-score of 4 was used for visualization.

Alt text: Bar plot showing expression dynamics of selected CAZyme classes in falcate and oval asexual spores across developmental stages and early maize leaf infection (1 dpi).

Figure M: Illustration of the detailed expression dynamics (log₂ fold change) of major CAZyme classes, along with their constituent families, across different developmental stages in falcate and oval conidia. As reference conditions served freshly harvested oval and falcate conidia for all time points tested for the corresponding spore type. Statistical significance was tested using a Wilcoxon test, p-values were adjusted with the Holm method. Significances provided denote differences between conidial types. The statistical significance indicates the differences between conidial types within each CAZyme class at the respective developmental stage.

Alt text: Box Plot showing detailed expression dynamics of major CAZyme classes and their families in falcate and oval conidia across developmental stages.

Figure N: Heatmap displaying gene expression of selected developmental genes in oval and falcate conidia after different developmental timepoints and early leaf infection. Differential expression was determined with a log2 fold change ≤ –1 or ≥ 1 and an adjusted p-value (padj) < 0.05. The heatmap was generated using iDEP.96, with red indicating up-regulated genes and blue indicating down-regulated genes. Hierarchical clustering was performed based on correlation distance with average linkage. Z-score normalization (centered by subtracting the mean) was applied, and a cut-off Z-score of 4 was used for visualization.

Alt text: Heatmap showing the expression profiles of selected genes relevant for fungal development in falcate and oval conidia, with color codes from red (up-regulated genes) to blue (down-regulated genes).

Figure O: Heatmap highlighting known secondary metabolite gene clusters associated with the biosynthesis of fusaridione A, depudecin, and phomopsins (A, B, E), which are induced during leaf infection (1dpi) in both *C. graminicola* spore types. Red color depicts the up-regulated genes, whereas blue represents the down-regulated genes. The heatmap was generated using iDEP.96, with red indicating up-regulated genes and blue indicating down-regulated genes. Hierarchical clustering was performed based on correlation distance with average linkage. Z-score normalization (centered by subtracting the mean) was applied, and a cut-off Z-score of 4 was used for visualization.

Alt text: Heatmap showing the expression profiles of selected secondary metabolite biosynthesis clusters in falcate and oval conidia, with color codes from red (up-regulated genes) to blue (down-regulated genes).

## Supplementary Table headers (File S1)

Table A: Description and mapping rates of RNA-seq samples. Wt in this table refers to CgM2 wild type like strain of *Colletotrichum graminicola* (M1.001). Reads were mapped to the *C. graminicola* genome version V5 with the mapping rate (in %) given in the last column. Mapping rates range from 93 to 98 % except for samples 11-16 (LF and LO), where mapping rates are much lower than for the other samples. The reason for the low mapping rates in samples 11-16 is because these samples are derived from maize plants infected with falcate or oval conidia, and the majority of reads from these samples map to the plant genome (Mapping to confirm this was done to the genome of *Zea mays*, acc. no. GCF_902167145.1, to which 89-91 % of reads from samples 11-16 mapped).

Table B: Pathway enrichment analysis performed for freshly harvested conidia and early maize leaf infection. Pathway enrichment analysis was performed using the GAGE method implemented in iDEP.96 (log2 fold change <=-1 or >=1, padj <0.05). Gene sets ranging from 5 to 2000 genes were included. Pathways were considered significant at an FDR cutoff of 0.05. Prior to enrichment analysis, genes with FDR ≥ 1 were excluded. Pathways with blue labelled up are up-regulated, whereas pathways with red labelled down are down-regulated in specified samples.

Table C: Oligonucleotides used in this study.

Table D: Effector genes showing the highest spore-type dependent specificity during early leaf interactions of *C. graminicola*. Evaluation based on the Top 100 regulated effector genes (Figure H File S1). Genes were included when the log2 fold change was >2 comparing early leaf infection samples of falcate with early leaf infection samples of oval conidia (LF_vs_LO) and padj was <0.05. Reference conditions as indicated.

