## Supplementary File S1 for "Spore type-specific gene expression profiles underlying development and leaf infection processes of *Colletotrichum graminicola*"

Supplementary Figures:

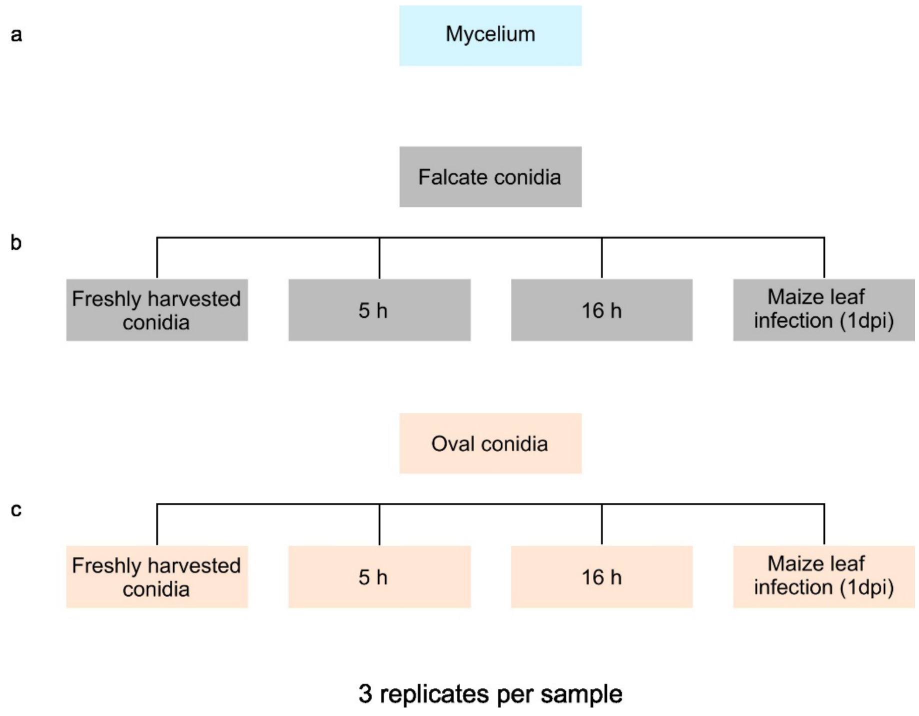


Figure A: Overview of RNA samples prepared. From each sample, including *C. graminicola* mycelium (a), falcate conidia (b), and oval conidia (c), 3 replicates were prepared. (b, c) Conidia samples were taken after different incubation timepoints including freshly harvested conidia, conidia incubated for the indicated timepoints under nutrient limiting conditions (23°C), and early maize leaf infection stages (1 dpi).

Alt text: Sample overview, with subfigures labelled from a to c, displaying the different samples used for the transcriptome analysis.

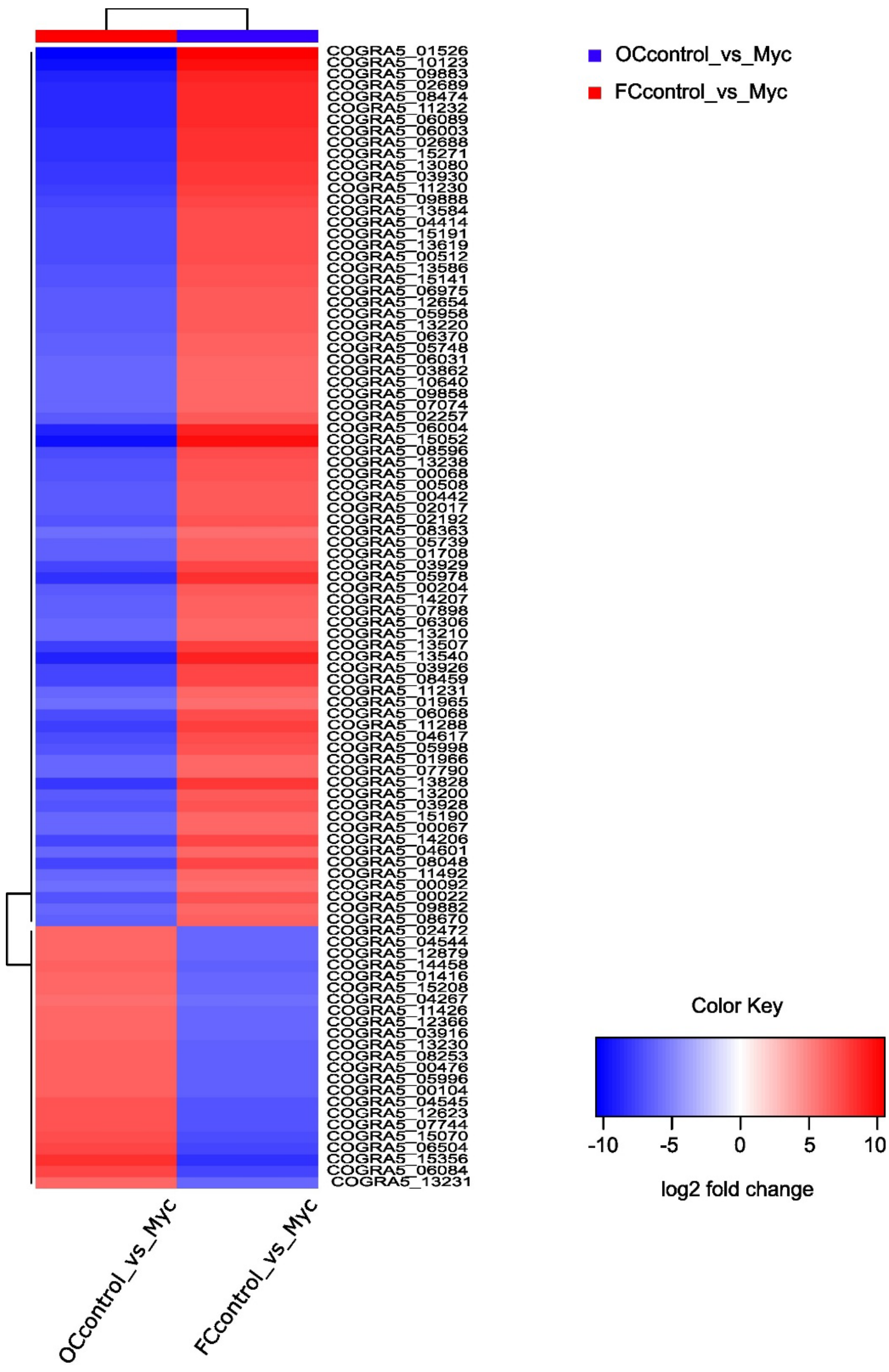


Figure B: Heatmap displaying the expression profiles of the 100 most variable genes in freshly harvested falcate and oval conidia. Red indicates up-regulation and blue indicates down-regulation of gene expression. The top 100 variable genes were identified using fold change values and were analyzed using iDEP.96 (Ge et al. 2018) where log2 fold change <=-1 or >=1, padj <0.05 was used. Hierarchical clustering was performed based on correlation distance with average linkage. Z-score normalization (centered by subtracting the mean) was applied, and a cut-off Z-score of 4 was used for analysis.

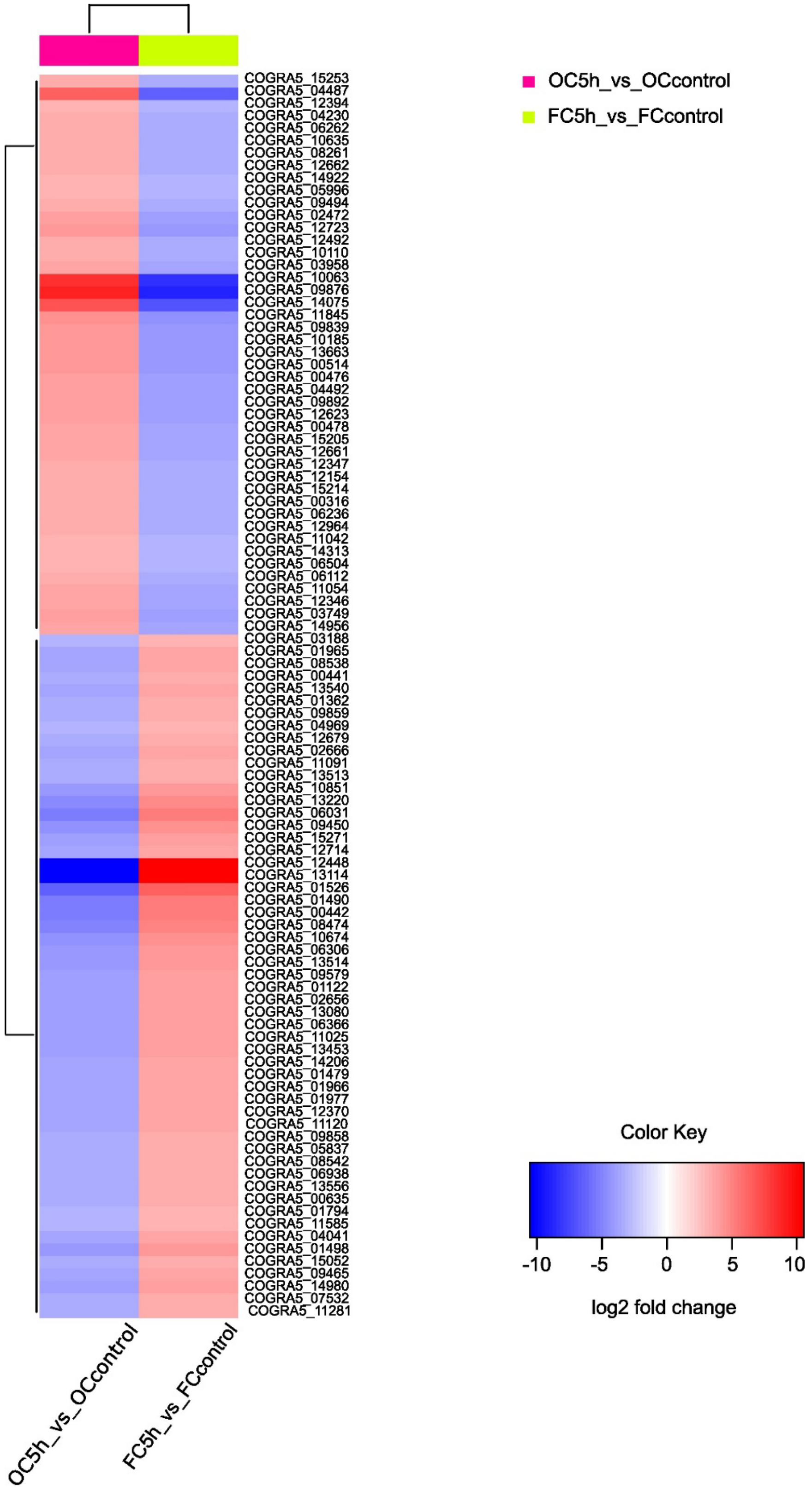
OC5h_vs_OCcontrol
FC5h_vs_FCcontrol
COGRA5_15253
COGRA5_04487
COGRA5_12394
COGRA5_04230
COGRA5_06262
COGRA5_10635
COGRA5_08261
COGRA5_12662
COGRA5_14922
COGRA5_05996
COGRA5_09494
COGRA5_02472
COGRA5_12723
COGRA5_12492
COGRA5_10110
COGRA5_03958
COGRA5_10063
COGRA5_09876
COGRA5_14075
COGRA5_11845
COGRA5_09839
COGRA5_10185
COGRA5_13663
COGRA5_00514
COGRA5_00476
COGRA5_04492
COGRA5_09892
COGRA5_12623
COGRA5_00478
COGRA5_15205
COGRA5_12661
COGRA5_12347
COGRA5_12154
COGRA5_15214
COGRA5_00316
COGRA5_06236
COGRA5_12964
COGRA5_11042
COGRA5_14313
COGRA5_06504
COGRA5_06112
COGRA5_11054
COGRA5_12346
COGRA5_03749
COGRA5_14956
COGRA5_03188
COGRA5_01965
COGRA5_08538
COGRA5_00441
COGRA5_13540
COGRA5_01362
COGRA5_09859
COGRA5_04969
COGRA5_12679
COGRA5_02666
COGRA5_11091
COGRA5_13513
COGRA5_10851
COGRA5_13220
COGRA5_06031
COGRA5_09450
COGRA5_15271
COGRA5_12714
COGRA5_12448
COGRA5_13114
COGRA5_01526
COGRA5_01490
COGRA5_00442
COGRA5_08474
COGRA5_10674
COGRA5_06306
COGRA5_13514
COGRA5_09579
COGRA5_01122
COGRA5_02656
COGRA5_13080
COGRA5_06366
COGRA5_11025
COGRA5_13453
COGRA5_14206
COGRA5_01479
COGRA5_01966
COGRA5_01977
COGRA5_12370
COGRA5_11120
COGRA5_09858
COGRA5_05837
COGRA5_08542
COGRA5_06938
COGRA5_13556
COGRA5_00635
COGRA5_01794
COGRA5_11585
COGRA5_04041
COGRA5_01498
COGRA5_15052
COGRA5_09465
COGRA5_14980
COGRA5_07532
COGRA5_11281
OC5h_vs_OCcontrol
FC5h_vs_FCcontrol
Color Key
-10
-5
0
5
10
log2 fold change

Figure C: 100 most variable genes in falcate and oval conidia following 5 h of incubation at nutrient limiting conditions depicted in a heatmap. Expression levels are color-coded, with red representing up-regulated and blue representing down-regulated genes. The top 100 variable genes were identified using fold change values and were analyzed using iDEP.96 (Ge et al. 2018) where log2 fold change <=-1 or >=1, padj <0.05 was used. Hierarchical clustering was performed based on correlation distance with average linkage. Z-score normalization (centered by subtracting the mean) was applied, and a cut-off Z-score of 4 was used for analysis.

Alt text: Heatmap showing the expression profiles of the most regulated genes (Top 100) in falcate and oval conidia after 5 h of cultivation in nutrient-limiting conditions, with color codes from red (up-regulated genes) to blue (down-regulated genes).

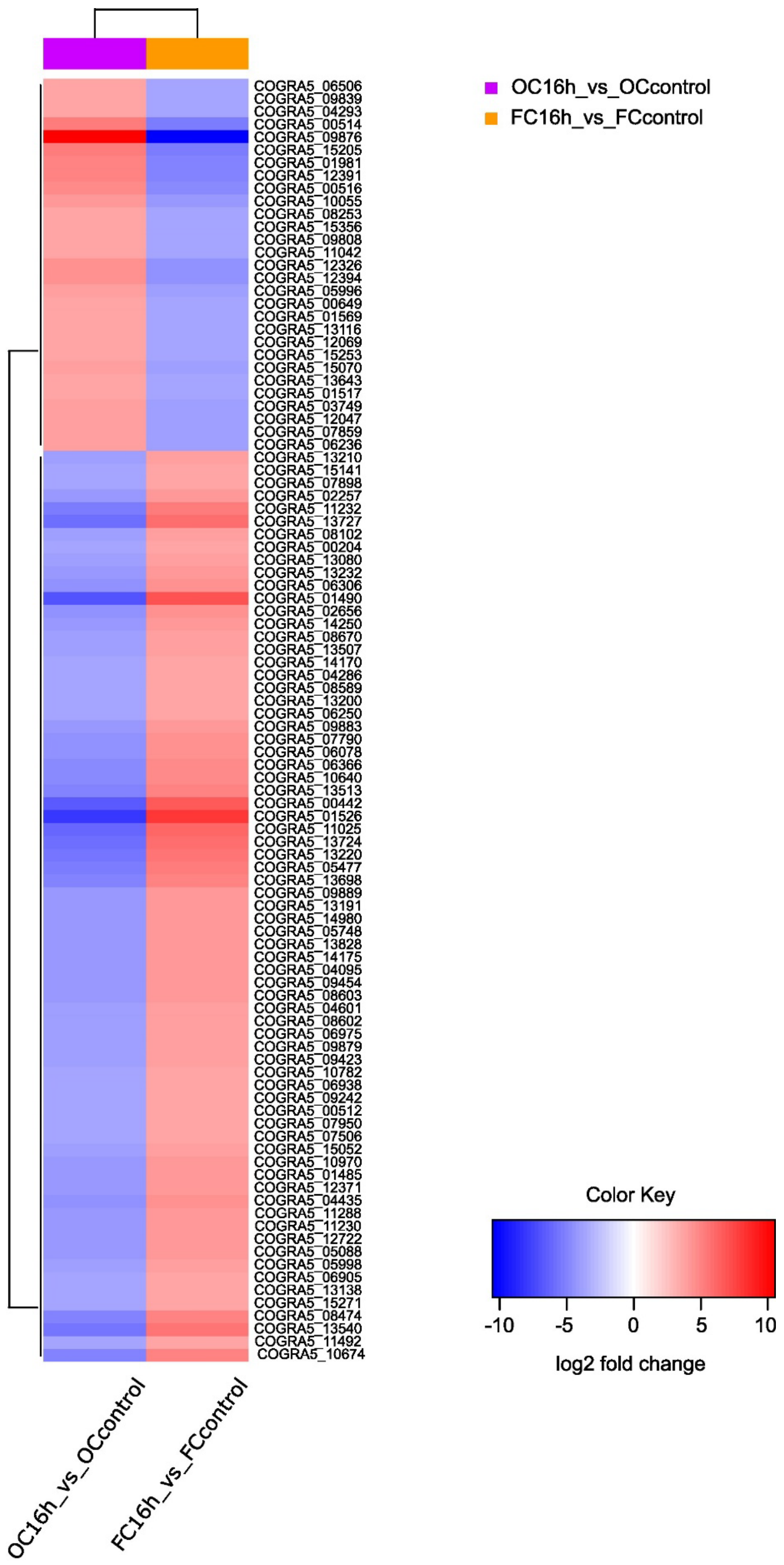
OC16h_vs_OCcontrol
FC16h_vs_FCcontrol
COGRA5_06506
COGRA5_09839
COGRA5_04293
COGRA5_00514
COGRA5_09876
COGRA5_15205
COGRA5_01981
COGRA5_12391
COGRA5_00516
COGRA5_10055
COGRA5_08253
COGRA5_15356
COGRA5_09808
COGRA5_11042
COGRA5_12326
COGRA5_12394
COGRA5_05996
COGRA5_00649
COGRA5_01569
COGRA5_13116
COGRA5_12069
COGRA5_15253
COGRA5_15070
COGRA5_13643
COGRA5_01517
COGRA5_03749
COGRA5_12047
COGRA5_07859
COGRA5_06236
COGRA5_13210
COGRA5_15141
COGRA5_07898
COGRA5_02257
COGRA5_11232
COGRA5_13727
COGRA5_08102
COGRA5_00204
COGRA5_13080
COGRA5_13232
COGRA5_06306
COGRA5_01490
COGRA5_02656
COGRA5_14250
COGRA5_08670
COGRA5_13507
COGRA5_14170
COGRA5_04286
COGRA5_08589
COGRA5_13200
COGRA5_06250
COGRA5_09883
COGRA5_07790
COGRA5_06078
COGRA5_06366
COGRA5_10640
COGRA5_13513
COGRA5_00442
COGRA5_01526
COGRA5_11025
COGRA5_13724
COGRA5_13220
COGRA5_05477
COGRA5_13698
COGRA5_09889
COGRA5_13191
COGRA5_14980
COGRA5_05748
COGRA5_13828
COGRA5_14175
COGRA5_04095
COGRA5_09454
COGRA5_08603
COGRA5_04601
COGRA5_08602
COGRA5_06975
COGRA5_09879
COGRA5_09423
COGRA5_10782
COGRA5_06938
COGRA5_09242
COGRA5_00512
COGRA5_07950
COGRA5_07506
COGRA5_15052
COGRA5_10970
COGRA5_01485
COGRA5_12371
COGRA5_04435
COGRA5_11288
COGRA5_11230
COGRA5_12722
COGRA5_05088
COGRA5_05998
COGRA5_06905
COGRA5_13138
COGRA5_15271
COGRA5_08474
COGRA5_13540
COGRA5_11492
COGRA5_10674
OC16h_vs_OCcontrol
FC16h_vs_FCcontrol
Color Key
-10
-5
0
5
10
log2 fold change

Figure D: Heatmap displaying 100 most variable genes in falcate and oval conidia following 16 h of incubation at nutrient limiting conditions. Up-regulated genes are depicted in red and down-regulated in blue. The top 100 variable genes were identified using fold change values and were analyzed using iDEP.96 (Ge et al. 2018) where log2 fold change <=-1 or >=1, padj <0.05 was used. Hierarchical clustering was performed based on correlation distance with average linkage. Z-score normalization (centered by subtracting the mean) was applied, and a cut-off Z-score of 4 was used for analysis.

Alt text: Heatmap showing the expression profiles of the most regulated genes (Top 100) in falcate and oval conidia after 16 h of cultivation in nutrient-limiting conditions, with color codes from red (up-regulated genes) to blue (down-regulated genes).

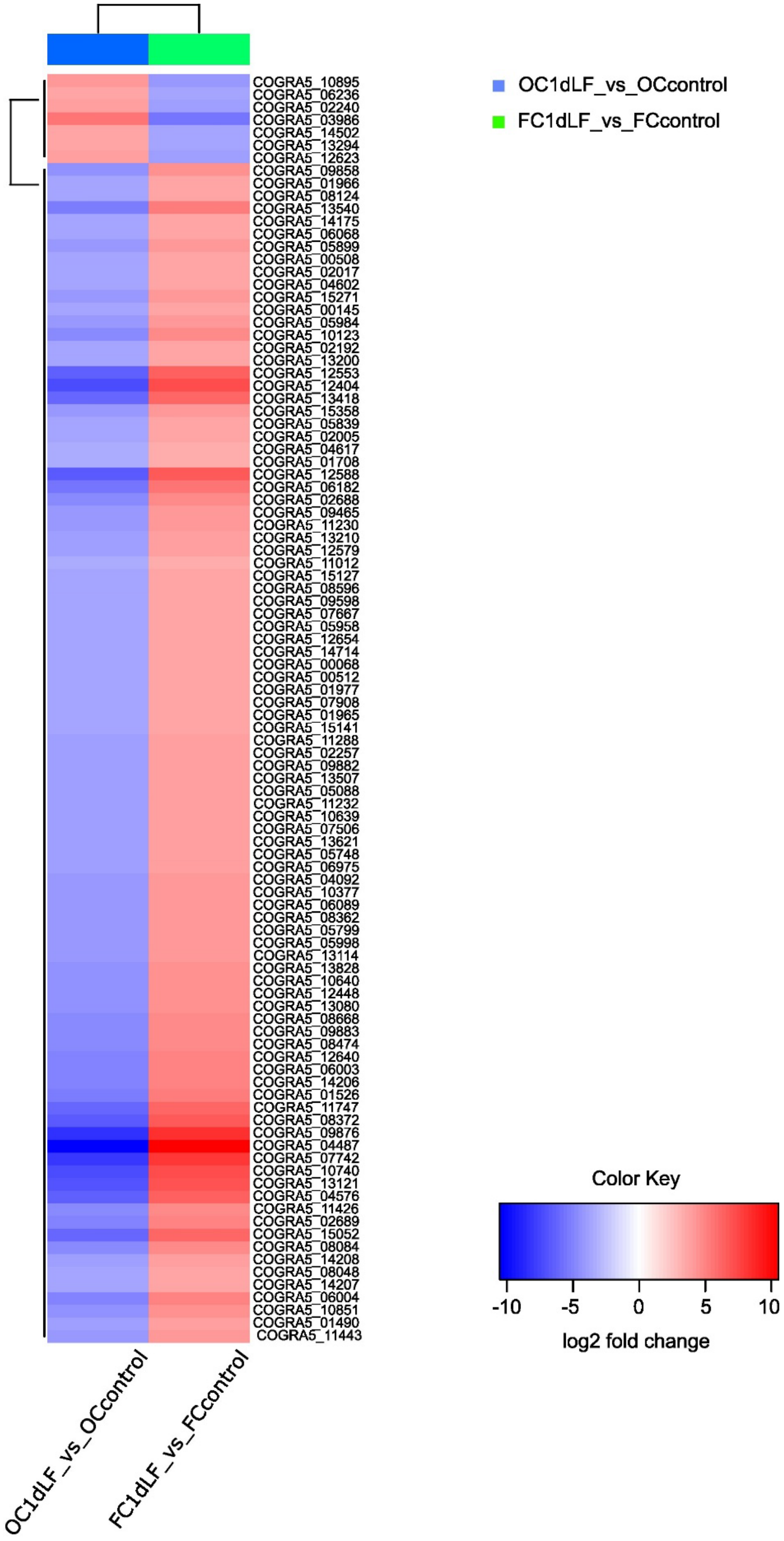

COGRA5_10895
COGRA5_06236
COGRA5_02240
COGRA5_03986
COGRA5_14502
COGRA5_13294
COGRA5_12623
COGRA5_09858
COGRA5_01966
COGRA5_08124
COGRA5_13540
COGRA5_14175
COGRA5_06068
COGRA5_05899
COGRA5_00508
COGRA5_02017
COGRA5_04602
COGRA5_15271
COGRA5_00145
COGRA5_05984
COGRA5_10123
COGRA5_02192
COGRA5_13200
COGRA5_12553
COGRA5_12404
COGRA5_13418
COGRA5_15358
COGRA5_05839
COGRA5_02005
COGRA5_04617
COGRA5_01708
COGRA5_12588
COGRA5_06182
COGRA5_02688
COGRA5_09465
COGRA5_11230
COGRA5_13210
COGRA5_12579
COGRA5_11012
COGRA5_15127
COGRA5_08596
COGRA5_09598
COGRA5_07667
COGRA5_05958
COGRA5_12654
COGRA5_14714
COGRA5_00068
COGRA5_00512
COGRA5_01977
COGRA5_07908
COGRA5_01965
COGRA5_15141
COGRA5_11288
COGRA5_02257
COGRA5_09882
COGRA5_13507
COGRA5_05088
COGRA5_11232
COGRA5_10639
COGRA5_07506
COGRA5_13621
COGRA5_05748
COGRA5_06975
COGRA5_04092
COGRA5_10377
COGRA5_06089
COGRA5_08362
COGRA5_05799
COGRA5_05998
COGRA5_13114
COGRA5_13828
COGRA5_10640
COGRA5_12448
COGRA5_13080
COGRA5_08668
COGRA5_09883
COGRA5_08474
COGRA5_12640
COGRA5_06003
COGRA5_14206
COGRA5_01526
COGRA5_11747
COGRA5_08372
COGRA5_09876
COGRA5_04487
COGRA5_07742
COGRA5_10740
COGRA5_13121
COGRA5_04576
COGRA5_11426
COGRA5_02689
COGRA5_15052
COGRA5_08084
COGRA5_14208
COGRA5_08048
COGRA5_14207
COGRA5_06004
COGRA5_10851
COGRA5_01490
COGRA5_11443
OC1dLF_vs_OCcontrol
FC1dLF_vs_FCcontrol
OC1dLF_vs_OCcontrol
FC1dLF_vs_FCcontrol
Color Key
-10
-5
0
5
10
log2 fold change

Figure E: Heatmap displaying 100 most variable genes in falcate and oval conidia harvested from maize leaves after 1 d of inoculation. The heatmap uses red and blue to denote up-regulation and down-regulation, respectively. The top 100 variable genes were identified using fold change values and were analyzed using iDEP.96 (Ge et al. 2018) where log2 fold change <=-1 or >=1, padj <0.05 was used. Hierarchical clustering was performed based on correlation distance with average linkage. Z-score normalization (centered by subtracting the mean) was applied, and a cut-off Z-score of 4 was used for analysis.

Alt text: Heatmap showing the expression profiles of the most regulated genes (Top 100) in falcate and oval conidia after 1 d of leaf inoculation, with color codes from red (up-regulated genes) to blue (down-regulated genes).

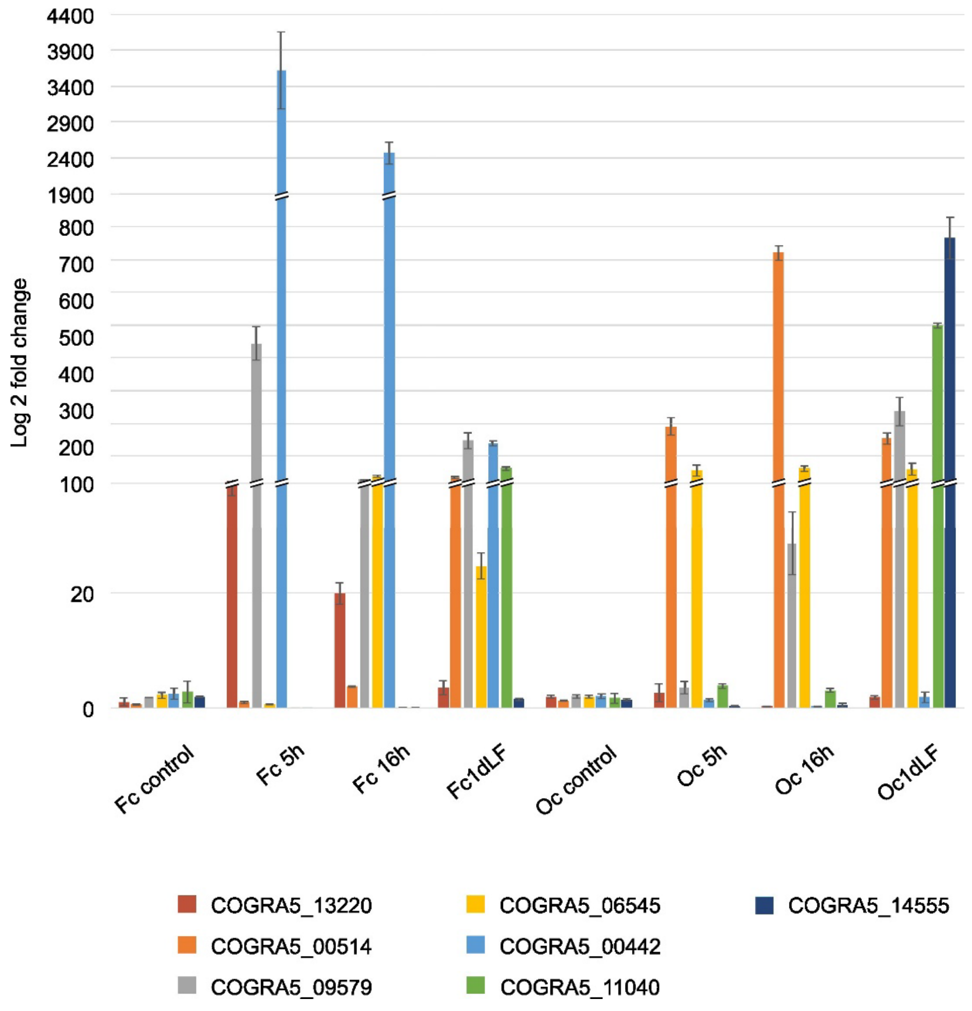


Figure F: Expression analysis of selected differentially expressed genes with qRT-PCR. RNA was taken from the same samples which were analyzed in the RNA-seq analysis. After DNA digestion, reverse transcription was used to generate cDNA, serving as template for the qRT-PCR reaction. Correct performance of the used oligonucleotides was tested beforehand. Base mean of the corresponding genes were: 388.88 (COGRA5_13220), 1152.99 (COGRA5_00514), 121.67 (COGRA5_09579), 260.96 (COGRA5_06545), 371.98 (COGRA5_00442), 227.90 (COGRA5_11040), 145.46 (COGRA5_14555).

Alt text: Bar chart showing qRT-PCR expression levels for selected differentially expressed genes.

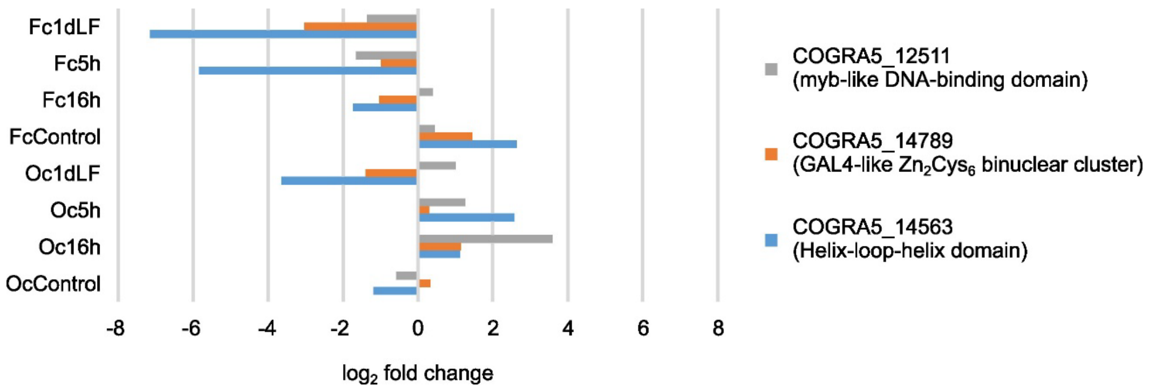


Figure G: Differentially expressed transcription factor-encoding genes showing increased expression in oval conidia relative to falcate conidia compared to the corresponding refence conditions across different developmental stages. As reference conditions for freshly harvested conidia (FcControl, OcControl) served mycelium, for all developmental and pathogenicity-related conditions (Fc5h, Fc16h, Fc1dLF, Oc5h, Oc16h, Oc1dLF) the corresponding freshly harvested conidia samples were chosen. The top 100 variable genes were identified using fold change values and were analyzed using iDEP.96 (Ge et al. 2018) where log2 fold change <=-1 or >=1, padj <0.05 was used. Hierarchical clustering was performed based on correlation distance with average linkage. Z-score normalization (centered by subtracting the mean) was applied, and a cut-off Z-score of 4 was used for analysis. Here different color bars represent different transcription factor genes.

Alt text: Bar chart showing differentially expressed transcription factor-encoding genes with higher expression in oval conidia than in falcate conidia.

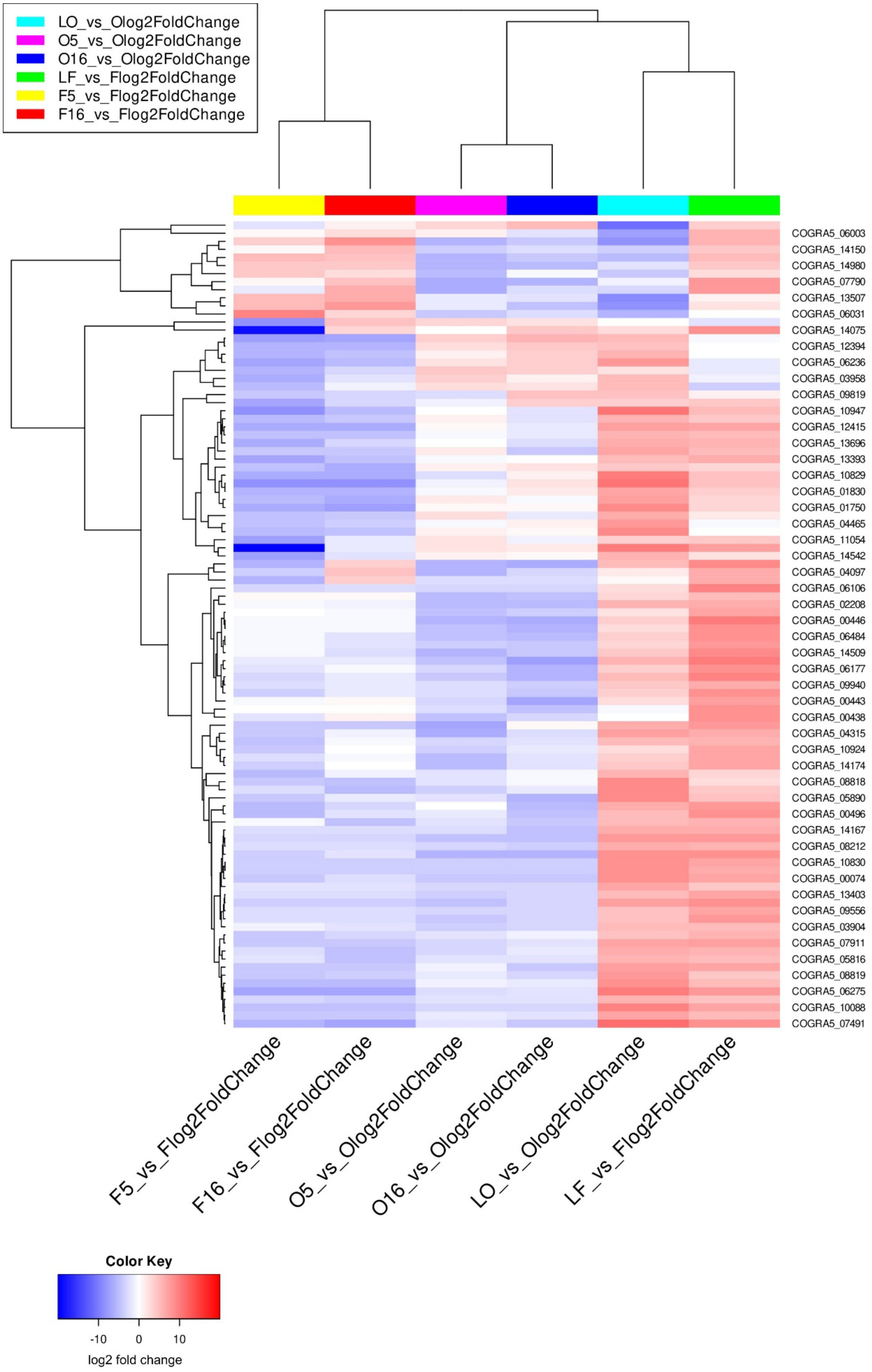
LO_vs_Olog2FoldChange
O5_vs_Olog2FoldChange
O16_vs_Olog2FoldChange
LF_vs_Flog2FoldChange
F5_vs_Flog2FoldChange
F16_vs_Flog2FoldChange
COGRA5_06003
COGRA5_14150
COGRA5_14980
COGRA5_07790
COGRA5_13507
COGRA5_06031
COGRA5_14075
COGRA5_12394
COGRA5_06236
COGRA5_03958
COGRA5_09819
COGRA5_10947
COGRA5_12415
COGRA5_13696
COGRA5_13393
COGRA5_10829
COGRA5_01830
COGRA5_01750
COGRA5_04465
COGRA5_11054
COGRA5_14542
COGRA5_04097
COGRA5_06106
COGRA5_02208
COGRA5_00446
COGRA5_06484
COGRA5_14509
COGRA5_06177
COGRA5_09940
COGRA5_00443
COGRA5_00438
COGRA5_04315
COGRA5_10924
COGRA5_14174
COGRA5_08818
COGRA5_05890
COGRA5_00496
COGRA5_14167
COGRA5_08212
COGRA5_10830
COGRA5_00074
COGRA5_13403
COGRA5_09556
COGRA5_03904
COGRA5_07911
COGRA5_05816
COGRA5_08819
COGRA5_06275
COGRA5_10088
COGRA5_07491
F5_vs_Flog2FoldChange
F16_vs_Flog2FoldChange
O5_vs_Olog2FoldChange
O16_vs_Olog2FoldChange
LO_vs_Olog2FoldChange
LF_vs_Flog2FoldChange
Color Key
-10
0
10
log2 fold change

Figure H: Heatmap to illustrate the 100 most variable expressed effector genes at different developmental stages in two conidia types. Heatmap was created using the iDEP.96 based on the corresponding log2 fold change values. Red color depicts the up-regulated genes, whereas blue represents the down-regulated genes. Hierarchical clustering was performed based on correlation distance with average linkage. Z-score normalization (centered by subtracting the mean) was applied, and a cut-off Z-score of 4 was used for visualization.

Alt text: Heatmap showing the expression profiles of the most regulated effector-encoding genes (Top 100) in falcate and oval conidia, with color codes from red (up-regulated genes) to blue (down-regulated genes).

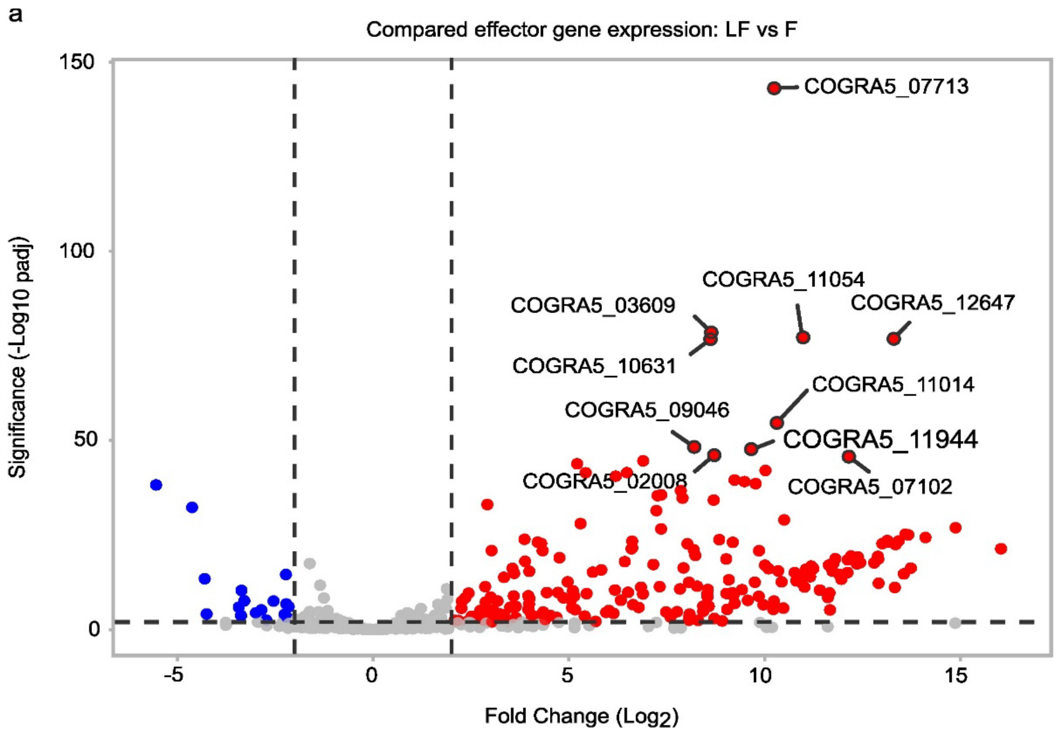


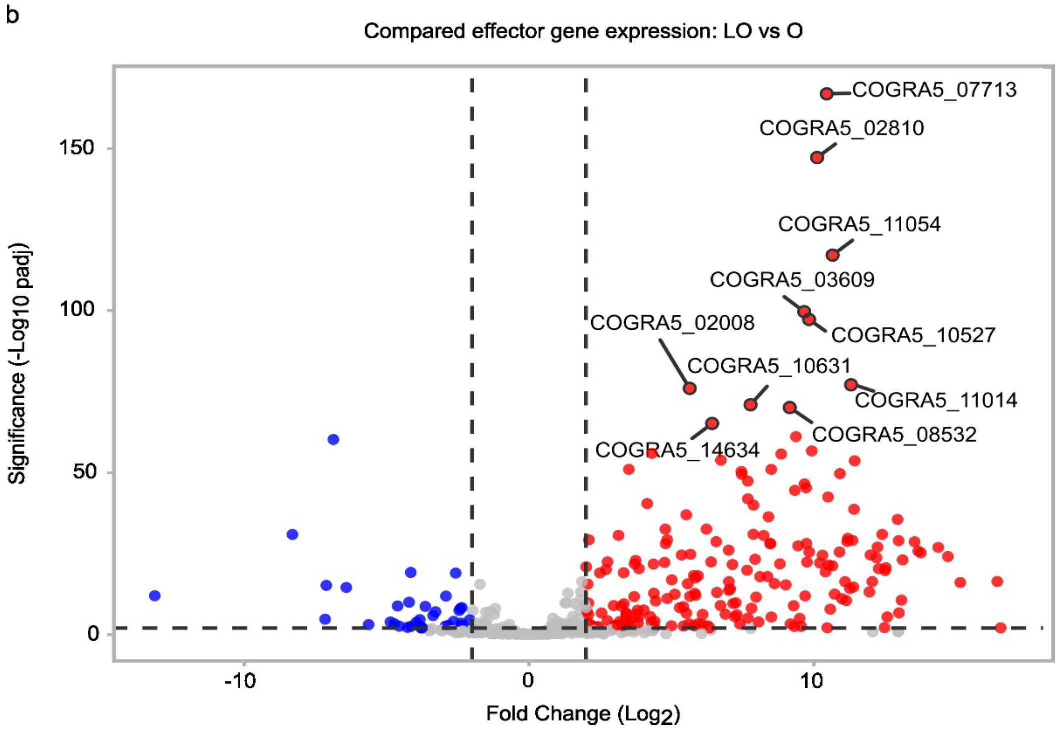


Figure I: Regulation of effector-encoding genes during early leaf infection. The volcano plots were generated based on the comparison of gene expression data of falcate conidia leaf infection vs freshly harvested falcate conidia (a) and oval conidia leaf infection vs freshly harvested oval conidia (b). VolcaNoseR settings were as followed: fold change threshold: -2 to 2; significant threshold: 2; use thresholds to annotate: changed and significant; criterion for ranking hits: Manhattan distance. Significantly regulated are indicated in red (up-regulated) and blue (down-regulated). Locus tag numbers of the 10 most relevant hits are indicated.

Alt text: Volcano plots showing regulation of effector-encoding genes during early maize leaf infection in oval and falcate conidia, with subfigures labelled from a to b, illustrating expression changes during early leaf infection with freshly harvested conidia as reference samples.

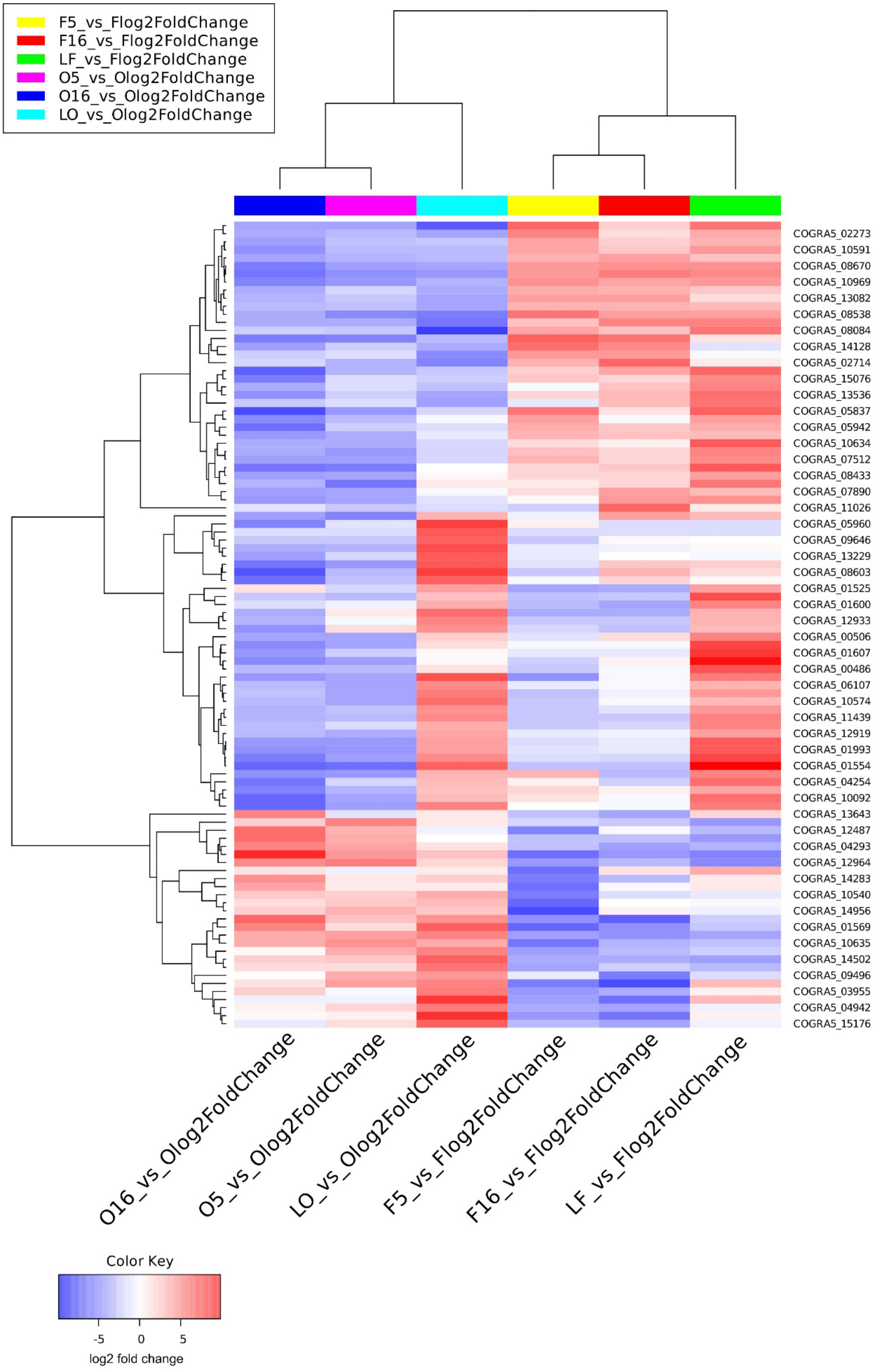

F5_vs_Flog2FoldChange
F16_vs_Flog2FoldChange
LF_vs_Flog2FoldChange
O5_vs_Olog2FoldChange
O16_vs_Olog2FoldChange
LO_vs_Olog2FoldChange
COGRA5_02273
COGRA5_10591
COGRA5_08670
COGRA5_10969
COGRA5_13082
COGRA5_08538
COGRA5_08084
COGRA5_14128
COGRA5_02714
COGRA5_15076
COGRA5_13536
COGRA5_05837
COGRA5_05942
COGRA5_10634
COGRA5_07512
COGRA5_08433
COGRA5_07890
COGRA5_11026
COGRA5_05960
COGRA5_09646
COGRA5_13229
COGRA5_08603
COGRA5_01525
COGRA5_01600
COGRA5_12933
COGRA5_00506
COGRA5_01607
COGRA5_00486
COGRA5_06107
COGRA5_10574
COGRA5_11439
COGRA5_12919
COGRA5_01993
COGRA5_01554
COGRA5_04254
COGRA5_10092
COGRA5_13643
COGRA5_12487
COGRA5_04293
COGRA5_12964
COGRA5_14283
COGRA5_10540
COGRA5_14956
COGRA5_01569
COGRA5_10635
COGRA5_14502
COGRA5_09496
COGRA5_03955
COGRA5_04942
COGRA5_15176
O16_vs_Olog2FoldChange
O5_vs_Olog2FoldChange
LO_vs_Olog2FoldChange
F5_vs_Flog2FoldChange
F16_vs_Flog2FoldChange
LF_vs_Flog2FoldChange
Color Key
-5
0
5
log2 fold change

Figure J: Heatmap to illustrate the 100 most variable transporter-encoding genes at different developmental stages in two conidia types. Heatmap was created using the iDEP.96 based on the corresponding log2 fold change values. Red color depicts the up-regulated genes, whereas blue represents the down-regulated genes. Hierarchical clustering was performed based on correlation distance with average linkage. Z-score normalization (centered by subtracting the mean) was applied, and a cut-off Z-score of 4 was used for visualization.

Alt text: Heatmap showing the expression profiles of the most regulated transporter-encoding genes (Top 100) in falcate and oval conidia, with color codes from red (up-regulated genes) to blue (down-regulated genes).

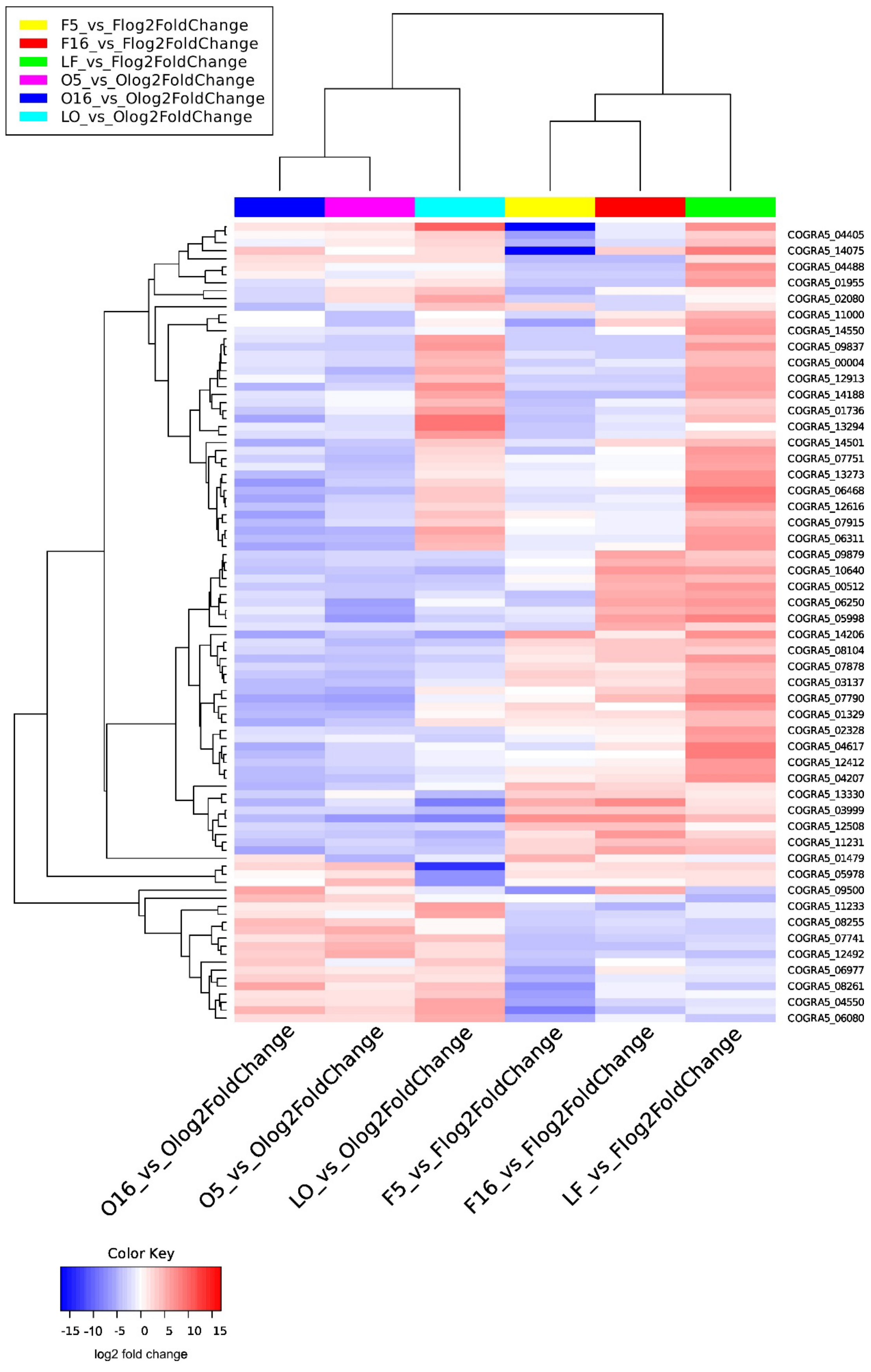
F5_vs_Flog2FoldChange
F16_vs_Flog2FoldChange
LF_vs_Flog2FoldChange
O5_vs_Olog2FoldChange
O16_vs_Olog2FoldChange
LO_vs_Olog2FoldChange
COGRA5_04405
COGRA5_14075
COGRA5_04488
COGRA5_01955
COGRA5_02080
COGRA5_11000
COGRA5_14550
COGRA5_09837
COGRA5_00004
COGRA5_12913
COGRA5_14188
COGRA5_01736
COGRA5_13294
COGRA5_14501
COGRA5_07751
COGRA5_13273
COGRA5_06468
COGRA5_12616
COGRA5_07915
COGRA5_06311
COGRA5_09879
COGRA5_10640
COGRA5_00512
COGRA5_06250
COGRA5_05998
COGRA5_14206
COGRA5_08104
COGRA5_07878
COGRA5_03137
COGRA5_07790
COGRA5_01329
COGRA5_02328
COGRA5_04617
COGRA5_12412
COGRA5_04207
COGRA5_13330
COGRA5_03999
COGRA5_12508
COGRA5_11231
COGRA5_01479
COGRA5_05978
COGRA5_09500
COGRA5_11233
COGRA5_08255
COGRA5_07741
COGRA5_12492
COGRA5_06977
COGRA5_08261
COGRA5_04550
COGRA5_06080
O16_vs_Olog2FoldChange
O5_vs_Olog2FoldChange
LO_vs_Olog2FoldChange
F5_vs_Flog2FoldChange
F16_vs_Flog2FoldChange
LF_vs_Flog2FoldChange
Color Key
-15 -10 -5 0 5 10 15
log2 fold change

Figure K: Heatmap to illustrate the 100 most variable genes encoding for Carbohydrate-Active Enzymes (CAZymes) at different developmental stages in two conidia types. Heatmap was created using the iDEP.96 based on the corresponding log2 fold change values. Red color depicts the up-regulated genes, whereas blue represents the down-regulated genes. Hierarchical clustering was performed based on correlation distance with average linkage. Z-score normalization (centered by subtracting the mean) was applied, and a cut-off Z-score of 4 was used for visualization.

Alt text: Heatmap showing the expression profiles of the most regulated CAZyme-encoding genes (Top 100) in falcate and oval conidia, with color codes from red (up-regulated genes) to blue (down-regulated genes).

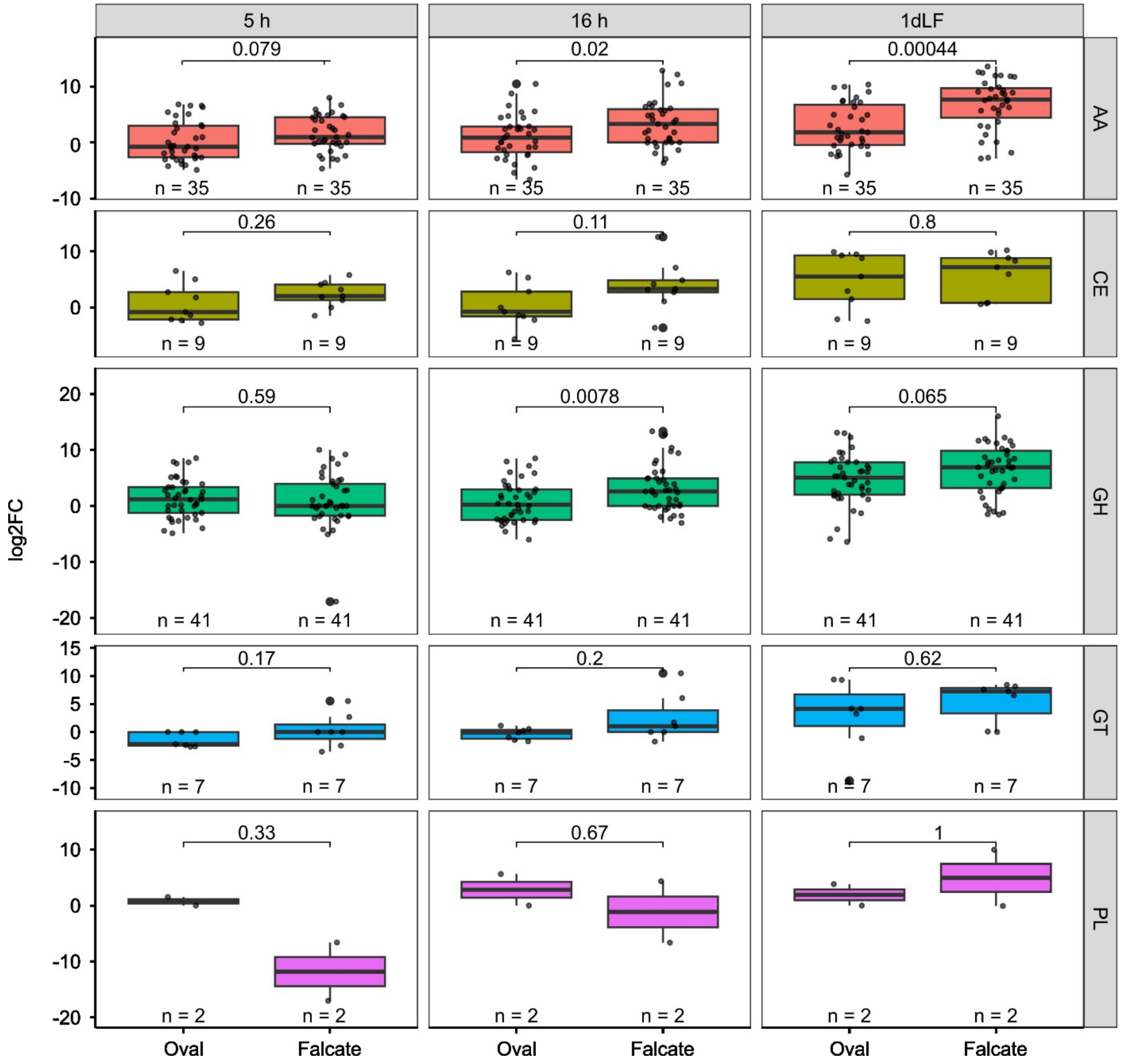


Figure L: Most variable differentially expressed Carbohydrate-Active Enzymes (CAZymes) in *C. graminicola* asexual spore types during leaf infection. Expression dynamics ($log_2$ fold change) of major CAZyme classes across different developmental stages and early maize leaf infection (1 dpi). As reference conditions served freshly harvested oval and falcate conidia for all time points tested for the corresponding spore type. Statistical significance was tested using a Wilcoxon test, p-values were adjusted with the Holm method. Significances displayed give differences between conidial types within each CAZyme class at the corresponding developmental stage, n = numbers of genes included. Identification of the top 100 variable genes was performed using iDEP.96

based on differentially expressed genes identified (DEseq). Hierarchical clustering was performed based on correlation distance with average linkage. Z-score normalization (centered by subtracting the mean) was applied, and a cut-off Z-score of 4 was used for visualization.

Alt text: Bar plot showing expression dynamics of selected CAZyme classes in falcate and oval asexual spores across developmental stages and early maize leaf infection (1 dpi).

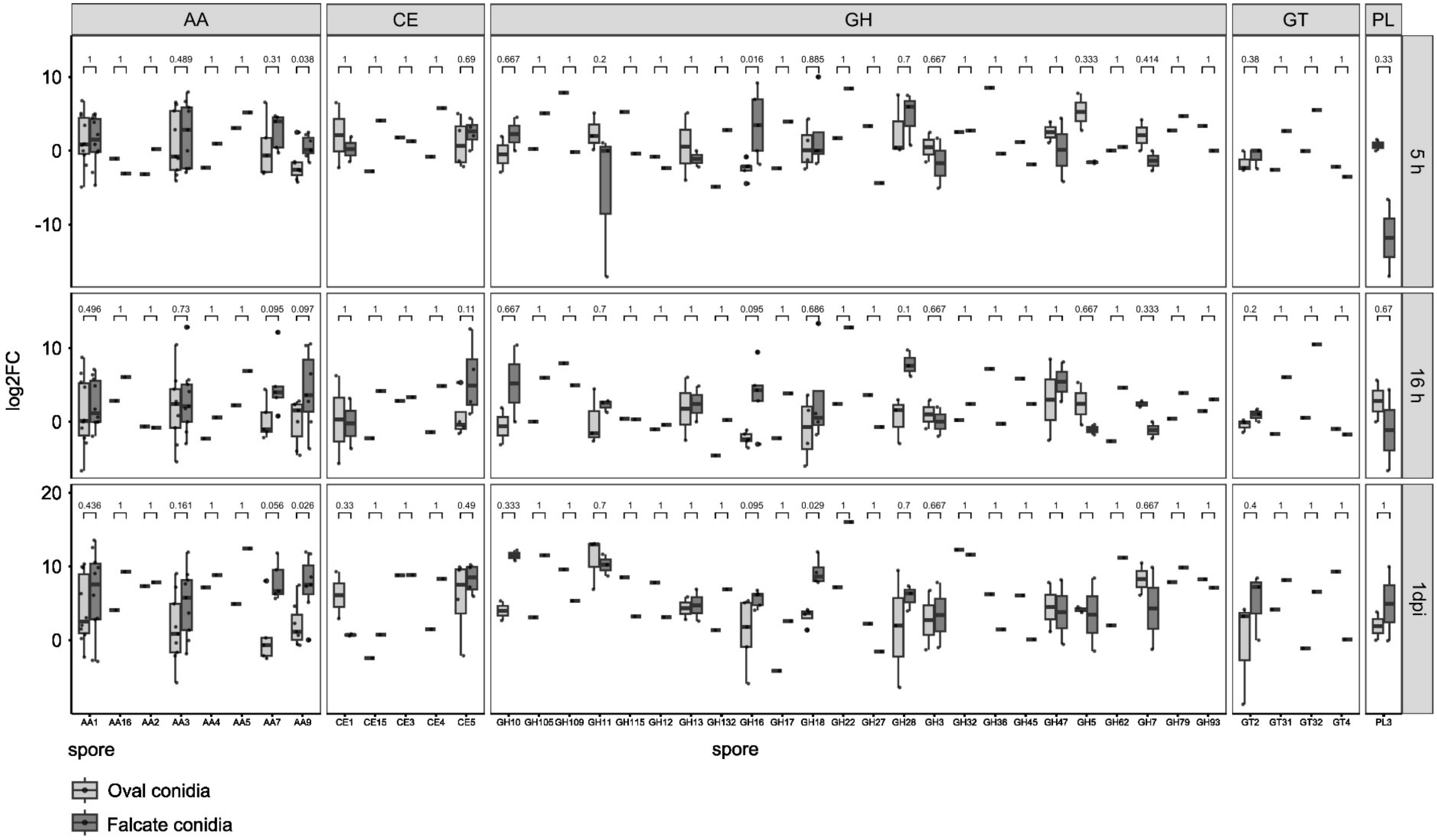


Figure M: Illustration of the detailed expression dynamics (log$_2$ fold change) of major CAZyme classes, along with their constituent families, across different developmental stages in falcate and oval conidia. As reference conditions served freshly harvested oval and falcate conidia for all time points tested for the corresponding spore type. Statistical significance was tested using a Wilcoxon test, p-values were adjusted with the Holm method. Significances provided denote differences between conidial types. The statistical significance indicates the differences between conidial types within each CAZyme class at the respective developmental stage.

Alt text: Box Plot showing detailed expression dynamics of major CAZyme classes and their families in falcate and oval conidia across developmental stages.

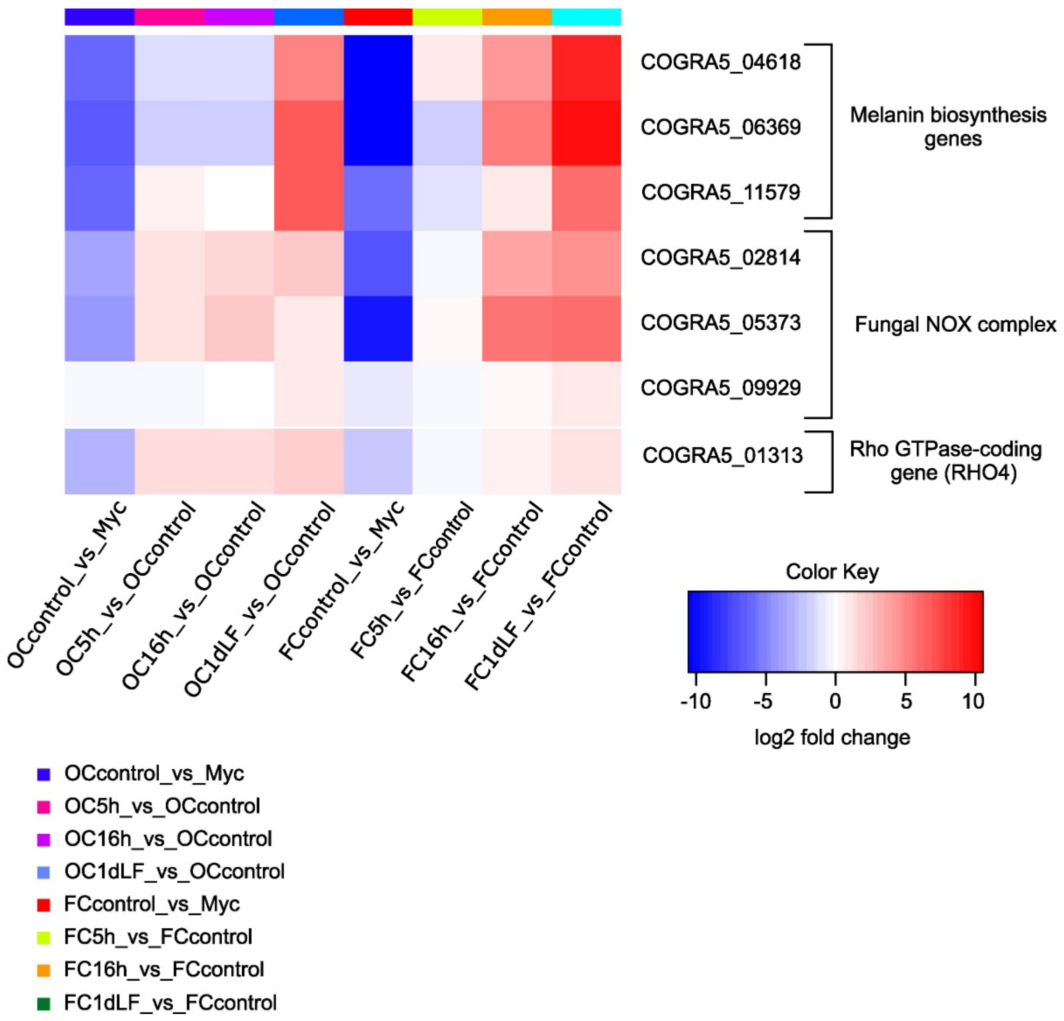


Figure N: Heatmap displaying gene expression of selected developmental genes in oval and falcate conidia after different developmental timepoints and early leaf infection. Differential expression was determined with a $\log_2$ fold change ≤ –1 or ≥ 1 and an adjusted p-value (padj) < 0.05. The heatmap was generated using iDEP.96, with red indicating up-regulated genes and blue indicating down-regulated genes. Hierarchical clustering was performed based on correlation distance with average linkage. Z-score normalization (centered by subtracting the mean) was applied, and a cut-off Z-score of 4 was used for visualization.

Alt text: Heatmap showing the expression profiles of selected genes relevant for fungal development in falcate and oval conidia, with color codes from red (up-regulated genes) to blue (down-regulated genes).

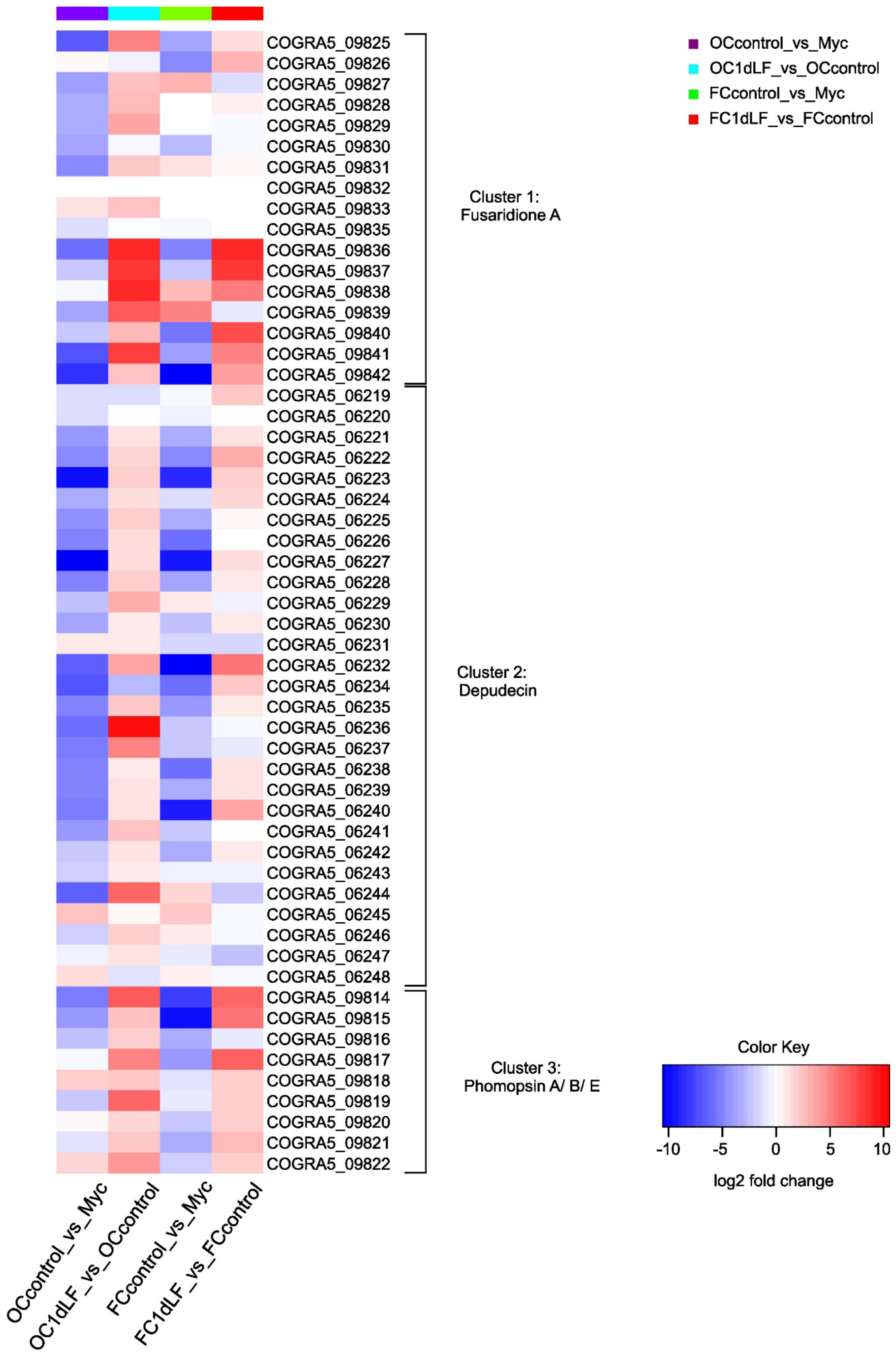

OCcontrol_vs_Myc
OC1dLF_vs_OCcontrol
FCcontrol_vs_Myc
FC1dLF_vs_FCcontrol
COGRA5_09825
COGRA5_09826
COGRA5_09827
COGRA5_09828
COGRA5_09829
COGRA5_09830
COGRA5_09831
COGRA5_09832
COGRA5_09833
COGRA5_09835
COGRA5_09836
COGRA5_09837
COGRA5_09838
COGRA5_09839
COGRA5_09840
COGRA5_09841
COGRA5_09842
COGRA5_06219
COGRA5_06220
COGRA5_06221
COGRA5_06222
COGRA5_06223
COGRA5_06224
COGRA5_06225
COGRA5_06226
COGRA5_06227
COGRA5_06228
COGRA5_06229
COGRA5_06230
COGRA5_06231
COGRA5_06232
COGRA5_06234
COGRA5_06235
COGRA5_06236
COGRA5_06237
COGRA5_06238
COGRA5_06239
COGRA5_06240
COGRA5_06241
COGRA5_06242
COGRA5_06243
COGRA5_06244
COGRA5_06245
COGRA5_06246
COGRA5_06247
COGRA5_06248
COGRA5_09814
COGRA5_09815
COGRA5_09816
COGRA5_09817
COGRA5_09818
COGRA5_09819
COGRA5_09820
COGRA5_09821
COGRA5_09822
Cluster 1: Fusaridione A
Cluster 2: Depudecin
Cluster 3: Phomopsin A/ B/ E
Color Key
-10
-5
0
5
10
log2 fold change

Figure O: Heatmap highlighting known secondary metabolite gene clusters associated with the biosynthesis of fusaridione A, depudecin, and phomopsins (A, B, E), which are induced during leaf infection (1dpi) in both *C. graminicola* spore types. Red color depicts the up-regulated genes, whereas blue represents the down-regulated genes. The heatmap was generated using iDEP.96, with red indicating up-regulated genes and blue indicating down-regulated genes. Hierarchical clustering was performed based on correlation distance with average linkage. Z-score normalization (centered by subtracting the mean) was applied, and a cut-off Z-score of 4 was used for visualization.

Alt text: Heatmap showing the expression profiles of selected secondary metabolite biosynthesis clusters in falcate and oval conidia, with color codes from red (up-regulated genes) to blue (down-regulated genes).

Supplementary Tables

Table A: Description and mapping rates of RNA-seq samples. Wt in this table refers to CgM2 wild type like strain of *Colletotrichum graminicola* (M1.001). Reads were mapped to the *C. graminicola* genome version V5 with the mapping rate (in %) given in the last column. Mapping rates range from 93 to 98 % except for samples 11-16 (LF and LO), where mapping rates are much lower than for the other samples. The reason for the low mapping rates in samples 11-16 is because these samples are derived from maize plants infected with falcate or oval conidia, and the majority of reads from these samples map to the plant genome (Mapping to confirm this was done to the genome of *Zea mays*, acc. no. GCF_902167145.1, to which 89-91 % of reads from samples 11-16 mapped).

| **Sample#** | **Sample description and abbreviation for downstream analysis** | **Number of reads** | **Mapping rate in %** |
|---|---|---:|---:|
| 2 | oval conidia after incubation for 5 h under nutrient limiting conditions (**O5**) | 24008487 | 97.7 |
| 3 | | 19847895 | 98.1 |
| 4 | | 20828345 | 97.7 |
| 5 | mycelium (**M**) | 20222305 | 97.7 |
| 6 | | 28996415 | 96.2 |
| 7 | | 22877299 | 97.0 |
| 8 | freshly harvested oval conidia (**O**) | 21225957 | 97.1 |
| 9 | | 27769533 | 97.2 |
| 10 | | 27985395 | 97.4 |
| 11 | leaf infection with falcate conidia 1dpi (**LF**) | 222299954 | 0.46 |
| 12 | | 178061742 | 0.94 |
| 13 | | 186226237 | 2.87 |
| 14 | leaf infection with oval conidia 1dpi (**LO**) | 207168903 | 0.74 |
| 15 | | 189665886 | 0.81 |
| 16 | | 158565123 | 0.69 |
| 17 | freshly harvested falcate conidia (**F**) | 21474610 | 94.6 |
| 18 | | 19909295 | 93.6 |

| | | | |
|---|---|---|---|
| 19 | | 24700766 | 94.6 |
| 20 | falcate conidia after incubation for 5 h under nutrient limiting conditions (**F5**) | 25167917 | 98.1 |
| 21 | | 24226438 | 97.8 |
| 22 | | 21012590 | 97.7 |
| 23 | falcate conidia after incubation for 16 h under nutrient limiting conditions (**F16**) | 22586828 | 98.1 |
| 24 | | 22042925 | 97.8 |
| 25 | | 26584963 | 97.6 |
| 26 | oval conidia after incubation for 16 h under nutrient limiting conditions (**O16**) | 26694901 | 98.3 |
| 27 | | 21464125 | 95.8 |
| 28 | | 26458053 | 94.7 |

Table B: Pathway enrichment analysis performed for freshly harvested conidia and early maize leaf infection. Pathway enrichment analysis was performed using the GAGE method implemented in iDEP.96 (log2 fold change <=-1 or >=1, padj <0.05). Gene sets ranging from 5 to 2000 genes were included. Pathways were considered significant at an FDR cutoff of 0.05. Prior to enrichment analysis, genes with FDR ≥ 1 were excluded. Pathways with blue labelled up are up-regulated, whereas pathways with red labelled down are down-regulated in specified samples.

| **Pathway** | **Fc control** | **Fc LF** | **Oc Control** | **Oc LF** |
|---|---|---|---|---|
| Secondary metabolic process | Down | Up | Down | Up |
| Mycotoxin biosynthetic process | Down | Up | Down | Up |
| Hydrolase activity, acting on glycosyl bonds | Down | Up | Down | Up |
| Carbohydrate metabolic process | Down | Up | Down | Up |
| Oxidoreductase activity | Down | Up | Down | Up |
| rRNA processing | Up | Down | Up | Down |
| Cellular response to stress | Up | Down | Up | Down |
| Cellular response to DNA damage stimulus | Up | Down | Up | Down |
| Nucleolus pathways | Up | Down | Up | Down |
| Protein dimerisation activity | Up | Down | Up | Down |

Table C: Oligonucleotides used in this study.

| Oligonucleotide | Sequence (5‘ to 3‘) |
|---|---|
| gapdh_fw | CCACCGACATGATCGGCAAC |
| gapdh_rv | GGGAGTAACCCCACTCGTTG |
| COGRA5_06545_fw | TGACACCTCCAACCAGCTTA |
| COGRA5_06545_rv | GTCCCAATCATCGAGGCAAG |
| COGRA5_14555_fw | GTCTACAACGAGCTGGCAAG |
| COGRA5_14555_rv | CTCTCCCGAACCTAACCCAG |
| COGRA5_11040_fw | CTGCCGACCTGAAATGGATG |
| COGRA5_11040_rv | CCCAAACGTACCTCCAAACC |
| COGRA5_00514_fw | GGGGTTCAACGGCAAGACTG |
| COGRA5_00514_rv | TGTCCTCGCACCAAGAATCTG |
| COGRA5_00442_fw | CTTACCAACTTCGCTTTGACAGG |
| COGRA5_00442_rv | GCCACCAATCGCATCGTTC |
| COGRA5_09579_fw | ACGAGACGTTGTGGGACGT |
| COGRA5_09579_rv | GAACCAGAGCGCTCTCGATC |
| COGRA5_13220_fw | CGACGTCTTGTACTTCGCGAC |
| COGRA5_13220_rv | GCAGCTCTTGAGGATGAGGC |

Table D: Effector genes showing the highest spore-type dependent specificity during early leaf interactions of *C. graminicola*. Evaluation based on the Top 100 regulated effector genes (Figure H File S1). Genes were included when the log2 fold change was >2 comparing early leaf infection samples of falcate with early leaf infection samples of oval conidia (LF_vs_LO) and padj was <0.05. Reference conditions as indicated.

| **locus_tag** | **base mean** | **LO_vs_O log2FoldChange** | **LF_vs_F log2FoldChange** | **LF_vs_LO log2FoldChange** | **effector type** | **reference** |
|---|---|---|---|---|---|---|
| COGRA5_06182 | 274,19 | -13,14809737 | 1,604954176 | 22,6848376 | Apoplastic effector | - |
| COGRA5_10559 | 172,60 | -7,162916069 | -2,210787695 | 6,992890832 | Apoplastic effector | (Kleemann et al. 2012, Oome and Van den Ackerveken 2014, Torres et al. 2016, Seidl and Van den Ackerveken 2019) |
| COGRA5_04185 | 23037,02 | -0,737109579 | 6,636901253 | 3,193825201 | Apoplastic effector | (Bettini et al. 2012, Eisermann et al. 2019) |
